# An Expanded Role for WRINKLED1 Metabolic Control Based on Combined Phylogenetic and Biochemical Analyses

**DOI:** 10.1101/2020.01.28.923292

**Authors:** Cathleen Kuczynski, Sean McCorkle, Jantana Keereetaweep, John Shanklin, Jorg Schwender

## Abstract

During triacylglycerol biosynthesis in developing oilseeds of *Arabidopsis thaliana*, fatty acid production is regulated by the seed-specific transcription factor WRINKLED1 (WRI1). WRI1 is known to directly stimulate the expression of fatty acid biosynthetic enzymes and a few targets in glycolysis. However, it remains unclear to what extent and how the conversion of sugars into fatty acid biosynthetic precursors in seeds is controlled by WRI1. Based on a previously reported DNA binding motif for WRI1, the ASML1/WRI1 (AW)- box, we developed a comparative genomics approach to search for conserved binding motifs in upstream regions of *Arabidopsis thaliana* protein-encoding genes and orthologous regions of 11 other Brassicaceae species. The AW-box was over-represented across orthologs for 915 *Arabidopsis thaliana* genes. Among these, 73 genes with functions in the biosynthesis of fatty acids and triacylglycerols and in glycolysis were enriched. For 90 AW-box sequences associated with these target genes, binding affinity to heterologously expressed *Arabidopsis thaliana* WRI1 protein was determined using Microscale Thermophoresis. Sites with low dissociation constants are preferentially located close to the transcriptional start site and are highly conserved between the 12 Brassicaceae species. Most of the associated genes were found to be co-expressed with WRI1 during seed development. When 46 automatically and manually curated genes containing conserved AW-sites with high binding affinity are mapped to central metabolism, a conserved regulatory blueprint emerges that infers concerted control of contiguous pathway sections in fatty acid biosynthesis and glycolysis. Among unexpectedly identified putative targets of WRI1 are plastidic fructokinase, phosphoglucose isomerase and several transcription factors.

**One sentence summary:** A combined comparative genomics and in-vitro DNA binding assay approach was used to identify conserved binding sites for the WRINKLED1 transcription factor in central metabolism and lipid biosynthesis.

## INTRODUCTION

Triacylglycerols (TAG), also known as vegetable oils, are an energy dense resource produced by many plants and stored in seeds and other plant organs. Plant oils are important for human nutrition as well as renewable biomaterials and fuels (Chapman and Ohlrogge, 2012). During seed development in oilseed species such as *Arabidopsis thaliana*, TAG is synthesized and accumulated at high rates (Neuhaus and Emes, 2000). Within the developing embryo, sugar supplies (sucrose) provided by maternal tissues are converted by conventional pathways of sugar catabolism into energy cofactors and pyruvate, which is the carbon precursor for chloroplast localized fatty acid synthesis (FAS) (Ruuska et al., 2002; O’Grady et al., 2012). WRINKLED1 (WRI1), a transcriptional regulator of the APETALA2/ethylene-responsive element-binding protein (AP2/EREBP) family has been characterized as a seed-specific transcription factor with control over FAS during the synthesis of TAG in developing oilseeds (Focks and Benning, 1998; Cernac and Benning, 2004; Masaki et al., 2005). WRI1 orthologs have been identified and characterized in a variety of plant species (Kong et al., 2019). Maeo *et al*. (2009) identified a consensus for WRI1 binding sites in upstream regions of WRI1 target genes, designated ASML1/WRI1 (AW)-box. In several cases, specific DNA binding by recombinantly expressed WRI1 to AW-box sites has been shown by Electro Mobility Shift Assays (Maeo et al., 2009; Fukuda et al., 2013; Li et al., 2015). In addition, we recently demonstrated the use of Microscale Thermophoresis (MST) for *in-vitro* quantification of binding affinity (dissociation constants) between recombinantly expressed WRI1 and DNA fragments (Liu et al., 2019). The MST approach gives opportunity to quantitatively characterize the DNA binding affinity of WRI1 in a medium throughput manner.

Since there is good evidence for sucrose synthase, the entry point of sugars, as well as plastidic pyruvate kinase and subsequent steps of FAS to be under direct control of WRI1 (Baud et al., 2007; Baud et al., 2009; Maeo et al., 2009), it seems likely that additional enzyme steps in-between sucrose cleavage and FAS are direct targets of WRI1. Indeed, activities of hexokinase and pyrophosphate dependent phosphofructokinase (PFP), aldolase, phosphoglycerate mutase and enolase were found to be substantially reduced in developing seeds of WRI1 KO mutants (Focks and Benning, 1998; Baud and Graham, 2006), i.e. might also be under direct control of WRI1. Other studies point to plastidic phosphoglycerate mutase and enolase to be direct targets of WRI1 (Baud et al., 2007; Baud and Lepiniec, 2009). In addition, under conditions where transient overexpression of WRI1 homologs from different plant species resulted in induced oil accumulation in *Nicotiana benthamiana* leaves (Grimberg et al., 2015), transcripts of sucrose synthase, PFP, enzyme steps in lower glycolysis between 3-phosphoglyceric acid (3PGA) and pyruvate and of the phosphoenolpyruvate/phosphate translocator of the chloroplast envelope were found to be upregulated (Grimberg et al., 2015). Altogether, WRI1 is widely understood as a “master regulator” for the conversion of sucrose to fatty acids in developing seeds and other oil accumulating tissues (Baud and Lepiniec, 2008; Chapman and Ohlrogge, 2012), but current knowledge on direct gene targets of WRI1, specifically with regards to the conversion of sucrose to pyruvate, is sparse. Given that glycolysis, the oxidative pentose phosphate pathway (OPPP) and the Ribulose 1,5-bisphosphate carboxylase/oxygenase shunt (O’Grady et al., 2012) constitute a ramified network of sugar catabolism in developing seeds with distinct enzyme isoforms in part being co-localized in the cytosol and the chloroplast compartments, it is of interest to better resolve which specific sections of sugar catabolism might be targeted by WRI1.

Here we report the results of a genome wide search for conservation of the AW-box in 12 species of the mustard family (Brassicaceae), including *A. thaliana* as the reference organism. Among *A. thaliana* genes for which the AW-box was over-represented across orthologous upstream regions (OURs), genes of glycolysis, FAS, TAG biosynthesis as well as transcription factors related to oil synthesis were found to be significantly enriched. Experimental validation of these targets by determination of *in-vitro* DNA binding activity to WRI1 protein and analysis of conservation of these sites across species reveals a conserved metabolic blueprint, giving insight on how WRI1 orchestrates central metabolism during seed oil biosynthesis.

## RESULTS

### The AW-box is enriched in -1 to 500 bp upstream regions among fatty acid biosynthetic genes in *A. thaliana*

Maeo *et al*. (2009) reported the AW-box (5’-CNTNG(N)_7_CG-3’; N = A, T, C or G) to be present in promoter regions upstream the ATG start codon and close to the transcriptional start site (TSS) in 19 out of 46 searched FAS genes. This suggests that the AW-box is statistically enriched in upstream regions of genes of the FAS pathway. We therefore quantitatively evaluated the presence of the AW-box in *A. thaliana* gene models for the set of 52 *Arabidopsis* genes annotated by the ARALIP database to be involved in FAS (annotation ‘Fatty Acid Synthesis’) (Li-Beisson et al., 2013) relative to the genomic background (Table 1). AW-box pattern matches were collected within five different search windows of 500 bp size, defined relative to the position of the start codon as well as the stop codon of each gene model (Table 1). Any number of pattern matches per searched sequence was counted as a hit. For each of the searched genomic regions, the mean expectation value of hits per 52 sampled genes can be derived from the genome wide frequency of hits (Table 1). For the region -1 to -500 bp upstream ATG, 10.5 hits are expected per 52 sampled genes (Table 1), which is very similar to the expectation values of randomized controls (Table 1), confirming that the genome wide frequency of AW-box hits is according to random chance expectation (Table 1). In contrast to the mean expectation of 10.5 hits, 30 hits were found for the -1 to - 500 bp upstream region of the 52 FAS genes, which means the AW-box is substantially enriched above the genomic background (hypergeometric *p*-value 3 x 10^-9^, Table 1). For the other genomic regions that were probed, the number of hits for sampling the 52 FAS genes was always very close to the mean expectation number, *i.e.* the AW-box was not over-represented relative to the background expectation (Table 1). Since for the -1 to -500 bp upstream region the mean expectation (10.5) amounts to 35 % of the detected number of hits (30), a substantial fraction of the 30 observed hits could be false positive. Given this level of anticipated spurious AW-box hits we supposed that identification of WRI1 targets could benefit from a phylogenetic footprinting approach as well as from testing individual binding sites in a medium throughput fashion by a quantitative *in-vitro* DNA binding assay for *At*WRI1 (Liu et al., 2019). Fig. 1 summarizes a workflow developed in this study which lead to the identification of numerous highly conserved AW-sites with *in-vitro* binding affinity to WRI1 protein, and therefore identifies likely WRI1 genes targets in *A. thaliana*, mostly within central metabolism.

**Figure 1.**
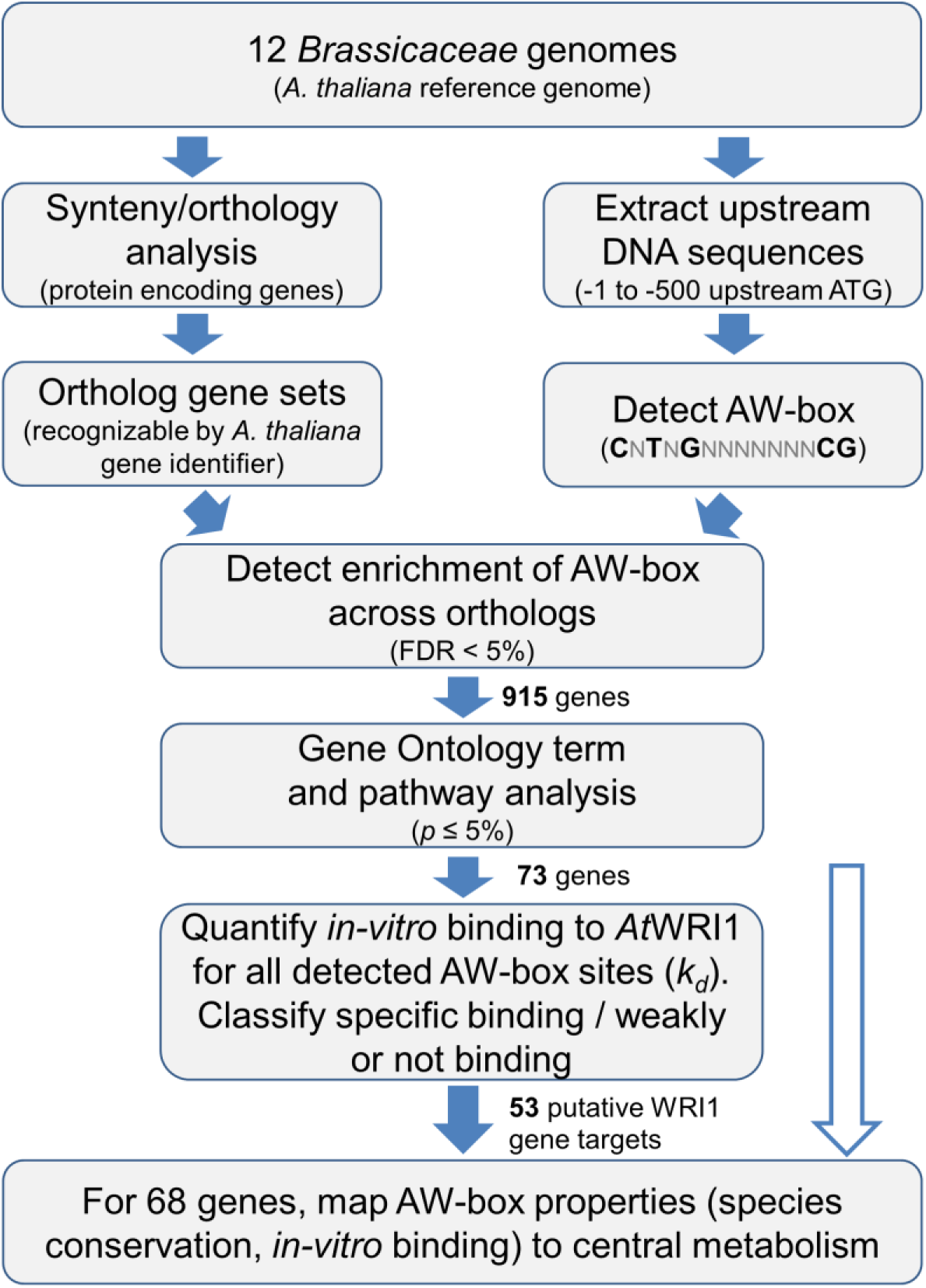
Workflow describing the main strategy developed in this study for detection of conserved and *in-vitro* binding AW-box sites. The white arrow indicates manual curated analysis of AW-box conservation and *in-vitro* binding for 21 additional genes of interest, as mentioned in the text.

**Table 1.** Enrichment of the AW-box motif in genomic features of genes encoding for fatty acid biosynthesis in *Arabidopsis thaliana*. For all protein encoding genes, regions relative to the start or stop codon were searched for the AW-box motif. Presence of the AW-box (AW-box hits) within the set of 52 FAS genes was modeled by the hypergeometric distribution based on the genome wide background. More details and controls see Supplemental Table S2. content in Supplemental Table S2A. ^3^Sequences were randomly shuffled using the “fasta-shuffle-letters” tool from the MEME suite (Bailey et al., 2009) with default settings; ^4^pseudo-random 500 bp sequences generated with the FaBox online tool (http://users-birc.au.dk/palle/php/fabox/random_sequence_generator.php) (VILLESEN, 2007) were generated using the average GC content in the -1 to -500 bp upstream sequences as input (33%).

| genomic feature relative to start / stop codons | genome wide number of hits (frequency of hits [%]) | Mean expected number of hits for 52 draws [99.9% confidence range] | number of hits for 52 FAS genes | fold enrichment | hypergeometric *p*-value |
| --- | --- | --- | --- | --- | --- |
| 1 to 500 bp upstream start | 5540 (20.2)<sup>1</sup> | 10.5 [0, 20] | 30 | 2.85 | 3 x 10<sup>-9</sup> |
| 501 to 1000 bp upstream start | 5675 (20.7)<sup>1</sup> | 10.8 [1, 21] | 12 | 1.11 | 0.39 |
| 1001 to 1500 bp upstream start | 6188 (22.6)<sup>1</sup> | 11.7 [1, 21] | 12 | 1.02 | 0.52 |
| 500 bp downstream start | 11414 (41.6)<sup>2</sup> | 21.7 [10, 32] | 25 | 1.15 | 0.21 |
| 500 bp downstream stop | 4569 (16.7)<sup>2</sup> | 8.7 [0, 18] | 6 | 0.69 | 0.88 |
| Randomized controls for regions upstream start codon |  |  |  |  |  |
| 1 to 500 bp upstream start<sup>3</sup> | 5919 (21.6) | 11.2 [1, 21] | 9 | 0.80 | 0.82 |
| random sequences<sup>4</sup> | 5889 (21.5) | 11.2 [1, 21] | n/a | n/a | n/a |

<sup>1</sup> average GC content close to 33 %; <sup>2</sup>The frequency of hits is different from the upstream regions, which is explainable by differing GC contents in these regions. See additional controls for variability on GC

### Synteny analysis across 12 *Brassicaceae* genomes

To allow testing for enrichment of the AW-box across orthologous promoter regions, genomic information for 12 species within the Brassicaceae family was collected from public available sources, including *A. thaliana*, which was designated as the annotated reference organism (Table 2). To define sets of syntenic ortholog genes we first compared the *A. thaliana* genome to each of the other genomes in a pair-wise fashion by using the SynOrths tool (Cheng et al., 2012). For all species between 70 and 94 % of the protein encoding gene content was found in ortholog gene sets (Table 2), which is similar to the 68 to 92 % of *A. thaliana* gene orthologs reported for *Brassicaceae* genomes (Haudry et al., 2013) and confirms that *Brassicaceae* genomes tend to be highly syntenic. All pairwise orthology relations were further aggregated into 25545 sets of orthologous genes (Supplemental Fig. S1). Each ortholog gene set is identified by the *A. thaliana* gene locus within the set.

**Table 2.**
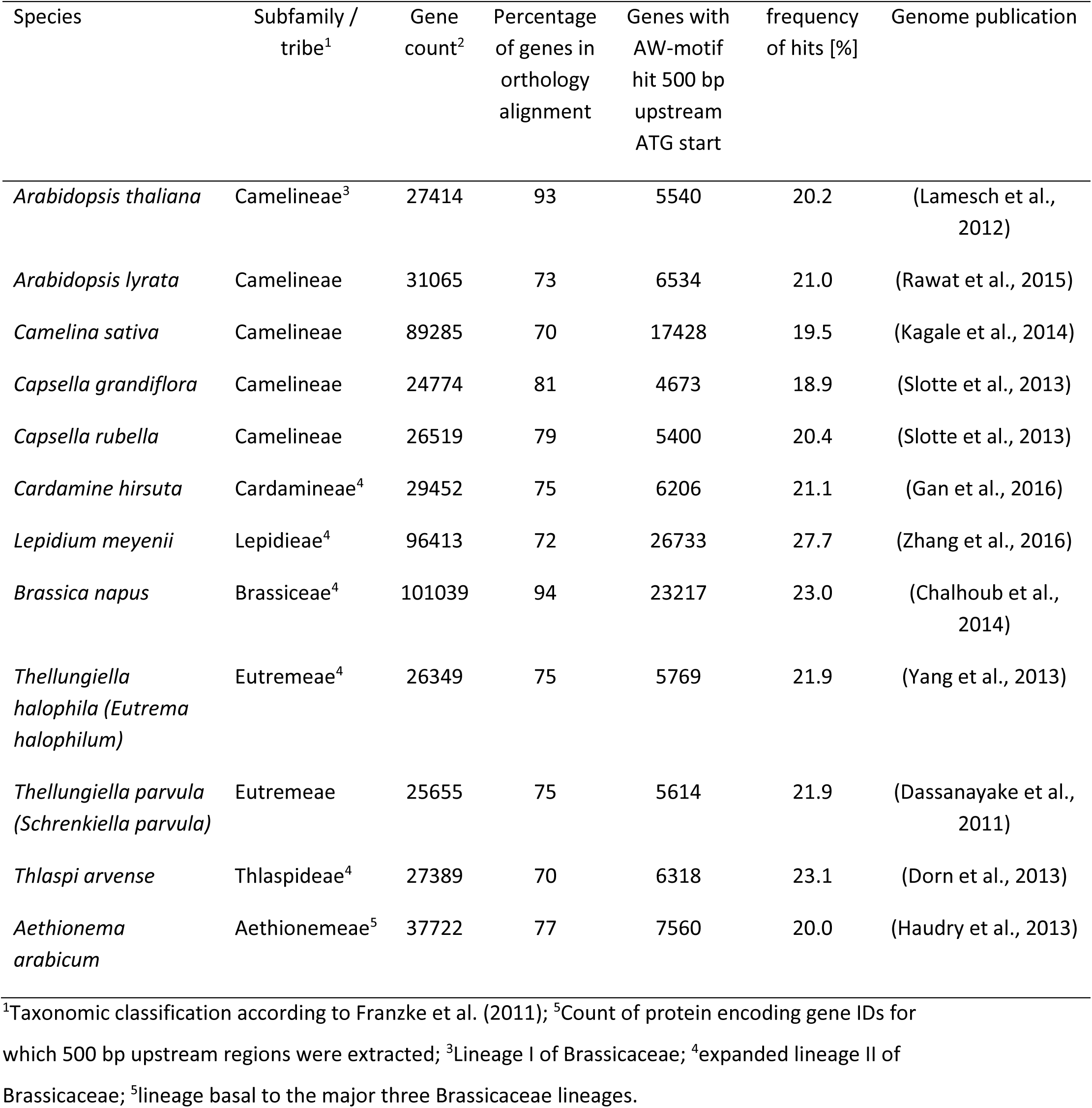
Species used in the phylogenetic foot printing approach and main results of genome mining for syntenic orthology relations and promoter regions. More details on data sources see Supplemental Table S1.

### The AW-box is significantly enriched across orthologous upstream regions of 915 *A. thaliana* genes and particularly for many genes of glycolysis and lipid biosynthesis

Next, the AW-box pattern was searched for in all 12 *Brassicaceae* species in 543076 genomic regions between -1 and -500 bp upstream the translational start of protein encoding genes (Table 2). Across all 12 searched genomes the AW-box was present for in-between 19.5 and 23 % of upstream sequences, which is similar to the frequency of 20.2 % that derives from the gene counts in in *A. thaliana* (Table 2). In assuming that true functional AW-box sites tend to be conserved across species, over-representation of AW-box hits among orthologous upstream regions (OURs) was tested for based on the cumulative hypergeometric probability function (Methods). The False Discovery Rate (FDR) was estimated based on re-determination of the hypergeometric *p*-values if the pattern search was repeated with random shuffled upstream sequences (Supplemental Fig. S2A). For a *p*-value threshold of 3.02 x 10^-4^ the empirical FDR was limited to 5 %. In this case 915 ortholog gene sets were judged to be significantly enriched with the AW box (Supplemental Fig. S2A). Accordingly, for 915 associated *A. thaliana* genes the AW-box is likely conserved, which is 6 times fewer than the 5540 AW-box hits that are obtained if only *A. thaliana* upstream regions are searched (Table 1). This indicates that the enrichment analysis can discriminate between conserved AW-box sites and spurious ones. Exploration of sequence alignments among AW-sites identified in the 915 ortholog gene sets revealed a substantial degree of conservation (Supplemental Fig. S3C-E). However, it is unclear if one can expect that the majority of the 915 discovered ortholog gene sets identify true functional WRI1 binding sites. We concluded that this is unlikely to be the case since repeating the AW-box enrichment analysis with base substitutions and permutations of the AW-box search pattern consistently resulted in close to 790 significance calls (Supplemental Fig. S2B).

The 915 *A. thaliana* gene identifiers for which the AW-box was found to be enriched across OURs were subjected to Gene Ontology (GO) term and pathway analysis (Table 3). Genes related to fatty acid, TAG and lipid synthesis were found to be significantly over-represented for all four classification systems tested (Table 3). The highest enrichment (12.9-fold) was found for category “Fatty Acid Synthesis” of the lipid pathways classification system ARALIP (Li-Beisson et al., 2013)(Table 3), the same gene set already considered for detection of the AW-box in the *A. thaliana* upstream region (Table 1). Also, 9 genes of ARALIP category “Triacylglycerol Biosynthesis” were identified (Table 3), which includes genes encoding enzyme functions in TAG biosynthesis as well as genes encoding for transcription factors. Furthermore, genes associated to glycolysis were significantly enriched in all classification systems except for ARALIP, which encompasses only lipid pathways (Table 3). Overall, over-representation of the AW-box was consistently found to be associated to lipid metabolism and glycolysis. The intersection of all enrichment gene sets in Table 3 identifies 73 *A. thaliana* gene loci. We postulated that a high proportion of these are direct WRI1 targets, which warrants detailed testing of binding specificity to WRI1 protein.

**Table 3.** Pathway and GO term enrichment for 915 *A. thaliana* genes for which the AW-box is significantly over-represented in upstream regions of ortholog gene sets. GO term analysis was performed using the DAVID online resource (Huang da et al., 2009), using the *A. thaliana* genes that identify the 25545 ortholog gene sets as background. Enrichment analysis for ARALIP lipid metabolic pathways (Li-Beisson et al., 2013), metabolic pathway categories of metabolic model Bna572 (Hay et al., 2014) and the MapMan classification system (Usadel et al., 2009) was performed as described in methods. All results with overrepresentation and a *p*-value < 0.05 are listed. For Mapman enrichment analysis only level 1 and 2 results are reported. See also Supplemental Table S3.

| classification system | Term / pathway | Expectation value | Gene count | Fold Enrichment | *p*-value* |
| --- | --- | --- | --- | --- | --- |
| Aralip | Fatty Acid Synthesis | 1.86 | 24 | 12.9 | 4 x 10<sup>-14</sup> |
| Aralip | Triacylglycerol Biosynthesis | 2.72 | 9 | 3.3 | 4 x 10<sup>-2</sup> |
| GO: Biological Process | glycolytic process (GO:0006096) | 2.69 | 17 | 6.3 | 6 x 10<sup>-6</sup> |
| GO: Biological Process | fatty acid biosynthetic process (GO:0006633) | 4.77 | 22 | 4.6 | 9 x 10<sup>-6</sup> |
| Bna572 | fatty acid synthesis | 3.12 | 21 | 6.7 | 2 x 10<sup>-10</sup> |
| Bna572 | glycolysis | 3.33 | 19 | 5.7 | 5 x 10<sup>-8</sup> |
| MapMan | lipid metabolism.FA synthesis and FA elongation (bin 11.1) | 3.51 | 24 | 6.8 | 1 x 10<sup>-11</sup> |
| MapMan | Glycolysis (bin 4) | 2.54 | 13 | 5.1 | 4 x 10<sup>-5</sup> |
| MapMan | TCA / org transformation (bin 8) | 2.72 | 10 | 3.7 | 1 x 10<sup>-2</sup> |
| MapMan | lipid metabolism (bin 11) | 15.0 | 36 | 2.4 | 5 x 10<sup>-5</sup> |

\*Adjusted *p*-value (Bonferroni correction).

### *in-vitro* binding assays of AW-box sequences from putative WRI1 target genes allow classification into binding and non-binding sites

The pathway and GO term enrichment analysis of Table 3 identified 73 *A. thaliana* genes. Within the 500 bp upstream region of these, because individual genes can have several AW boxes in this region, 95 AW-box sites are identified and for all of them binding affinities to recombinantly expressed AtWRI1 were determined based on MST (see methods). Since a few of the AW-box sites overlap, the 95 AW-boxes are represented by altogether 90 synthesized DNA fragments of 28 *bp* length for which dissociation constants (*k_d_*) were determined (Supplemental Table S4, measurements 1-86; 189-192). Fig. 2 shows that the resulting 90 *k_d_* values approximate a bimodal distribution which peaks for values close to 0 nM and for dissociation constants above 1000 nM which are grouped with non-binding DNA fragments. Taking advantage of the distinct bimodality of the distribution, DNA fragments were classified into specific binding and non-binding ones based on a threshold value of 200 nM, which designates the left peak to represent specific binding AW-sites (Fig. 2). While the choice of this specific threshold value is somewhat arbitrary, the exact value is not critical for our further analysis since the two maxima are clearly separated with only 9 *k_d_* values (10 % of the 90 values) spread between 200 and 1000 nM (Fig. 2). The 62 AW-box sites associated to specific binding DNA fragments (*K_d_* < 200 nM) have substantial overall sequence similarity that exceeds the five fully conserved bases of the AW-box pattern (Fig. 2, sequence logo *K_d_* < 200 nM). Notably, while the AW-box consensus originally was defined to be 14 nucleotide (nc) long (sequence logos in Fig. 2, positions 3 to 16), two nc positions outside the AW-box seem to be conserved among *in-vitro* WRI1 binding AW-sites (positions 17, 18). In contrast to the specific binding AW sites of the left peak of Fig. 2, no consensus beyond the 5 invariable bases of the AW-box is detectable for the 33 remaining sites of lower binding affinity (Fig. 2), suggesting that these instances are spurious pattern matches. To clearly demonstrate that the AW-box consensus alone does not reliably identify WRI1 binding sequences, we synthesized 10 additional 28 bp sized DNA fragments, the sequences of which were sampled from a pseudo-randomized DNA sequence background (33% G+C content), searching for an extended 28 nc AW-box pattern 5’-(N)_7_CNTNG(N)_7_CG(N)_7_-3’. In MST binding assays, only two *k_d_* values were just below 200 nM (150 nM, 185 nM), while 5 of the DNA fragments were judged to be non-binding (Supplemental Table S4, measurements 157-166).

**Figure 2.**
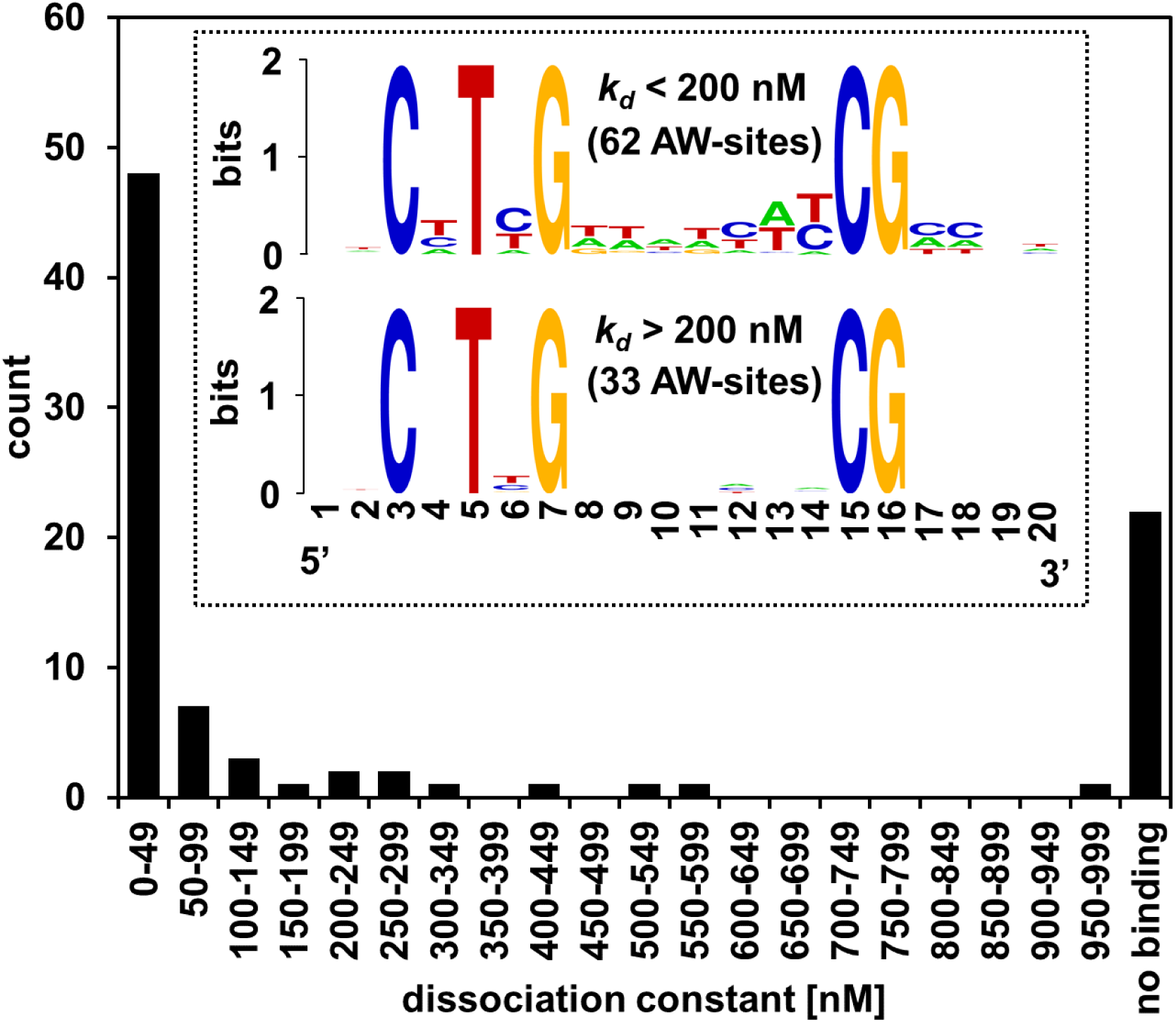
Distribution of DNA-WRI1 binding affinities for 90 AW-box sites found upstream of 73 likely WRI1 target genes in *A. thaliana*. The set of 73 *A. thaliana* genes was identified by pathway enrichment analysis (Table 3). Using Microscale thermophoresis, equilibrium dissociation constants (*K_d_*) were determined for 90 DNA targets, representing 95 putative WRI1 binding sites (AW-box sites) which are present upstream the genes. For sequence data and measurement values see Supplemental Table S4, measurements 1-86 and 189-192. Inset: Sequence logos of AW-box sites from measured DNA fragments, separated according to ranges of *k_d_* values. Sequence logos were generated with WebLogo (Crooks et al., 2004).

### AW-boxes that bind WRI1 tend to be phylogenetically conserved

For this study we defined *A. thaliana* as the reference species and sequence conservation was assessed by pairwise comparisons between *A. thaliana* AW-sites and sites of other species, with a stringent similarity cutoff (see methods). Pairwise conservation relations were aggregated to the gene and species level as demonstrated in Supplemental Fig. S3B. As an example, two AW-box sites positioned at -120 and -148 bp upstream the start codon of *A. thaliana* plastidic pyruvate kinase β_1_-subunit (*At*PK_p_β_1_) are shown in Fig. 3A. Both sites bind WRI1 *in-vitro* (Maeo et al., 2009)(Supplemental Table S4) and *in-vivo* functionality to drive WRI1 dependent gene expression has been thoroughly characterized for both sites (Maeo et al., 2009). The two AW-sites are conserved among 19 out of 22 OURs and among all of the 12 species of this study (Fig. 3A). Genes that miss a conserved site are PK_p_ isoforms in polyploid species (Fig. 3A). For both *At*PK_p_β_1_ AW-sites, sequence logos show full conservation at 13 base positions in all aligned sequences (Fig. 3B). To test if other AW-sites are similarly well conserved, the conservation analysis of Fig. 3A was applied to all AW-sites discovered in Fig. 2 by *in-vitro* binding assays (Supplemental Table S5). From the 73 *A. thaliana* genes with upstream AW-sites examined, 53 have at least one site classified to bind WRI1 specifically. Conservation was assessed by tracking only sites with specific binding activity (*k_d_* < 200 nM) and expressed as species conservation ratio, which is the number of species in which an *in-vitro* binding AW-site is conserved divided by 12 (total number of species). In result, in 79% of cases (42) the species conservation ratio is ≥ 0.8 (Fig. 3C). Many cases where the value of the species conservation ratio is below one can be explained by missing orthologs or the ortholog gene discovery process having missed to detect an ortholog in a species.

**Figure 3.**
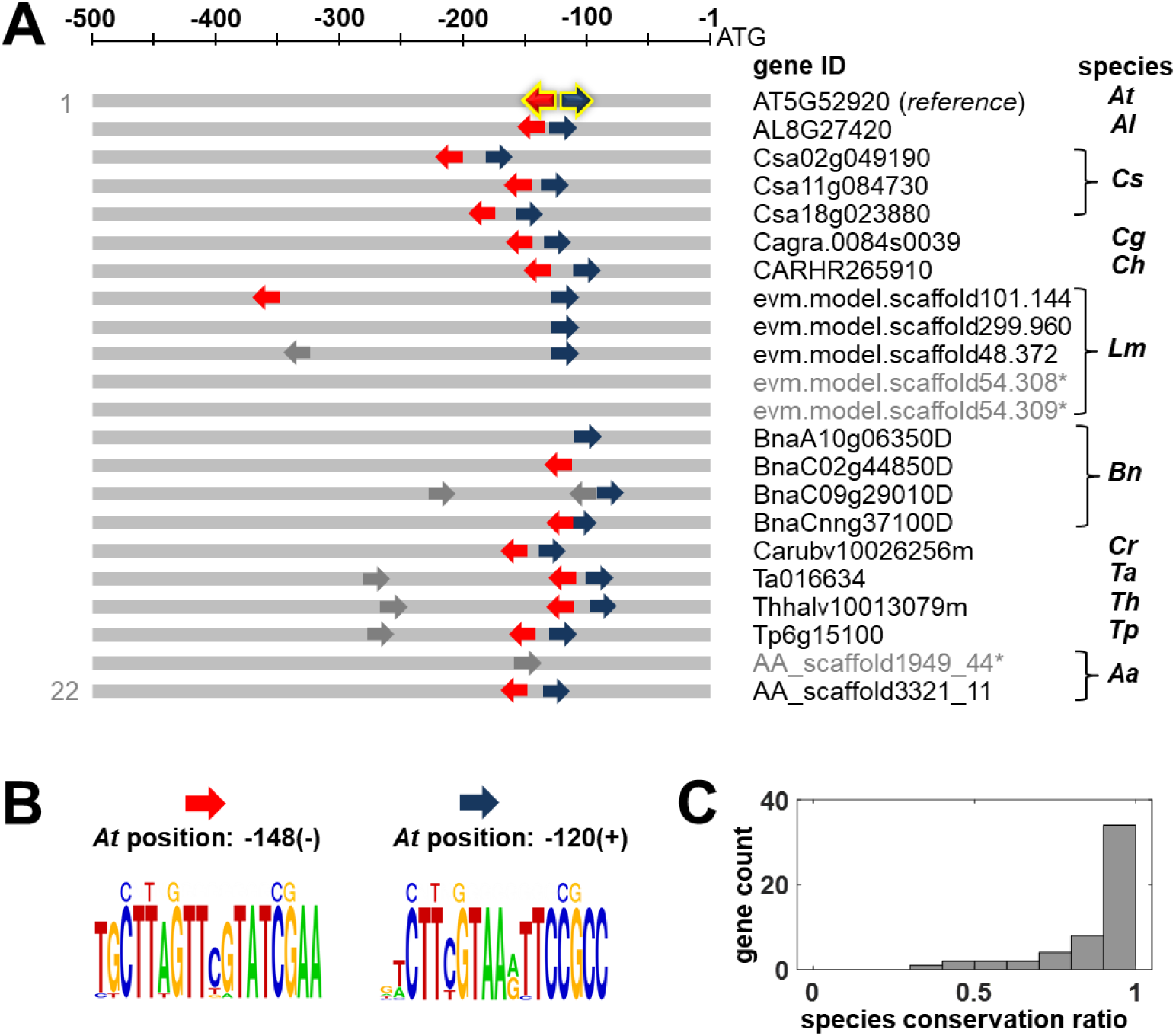
Enrichment and conservation of the AW-box motif across ortholog upstream regions (OURs). A, Relative position and orientation of two AW-box sites in the 500 bp upstream regions *A. thaliana* plastidic pyruvate kinase β_1_-subunit (AT5G52920) are conserved across orthologs. Although for three gene IDs (grey) no conserved AW sites were found, the two *A. thaliana* AW-sites are conserved across all 12 species of this study. B, Sequence logos of the two conserved AW-box sites. C, Species conservation ratio for 53 genes for which at least one AW-site binds WRI1 (*k_d_* < 200 nM, Fig. 2). Abbreviations: Aa, *Aethionema arabicum*; At, *Arabidopsis thaliana*; Al, *Arabidopsis lyrata*; Bn, *Brassica napus*; Cs, *Camelina sativa*; Cg, *Capsella grandiflora*; Cr, *Capsella rubella*; Ch, *Cardamine hirsuta*; Lm, *Lepidium meyenii*; Th, *Thellungiella halophila*; Tp, *Thellungiella parvula*; Ta, *Thlaspi arvense*.

If *A. thaliana* AW-box sites that are conserved and specifically bind WRI1 *in-vitro* are true functional sites, then one should expect that WRI1 binding affinity is also well preserved, i.e. that variation in AW-site sequences across orthologs should have only minor effects on WRI1 binding. We therefore selected 5 *A. thaliana* AW-sites (PK_p_β_1_ at upstream position -148; BCCP2, -29; KASI, -58; PGLM1; -231; PGLM2, -164) which are conserved in all 12 species of this study (Supplemental Table S5) to measure WRI1 affinities so we could compare them between aligned orthologous sequences (Fig. 4A-D). Due to the high degree of conservation, not all orthologous sequences were measured. All sequence variants of the 18 bp AW-box were evaluated. In all, 29 *k_d_* values that were measured for sequences in Fig. 4A-D fall well below the dissociation constant defined in Fig. 2 as a threshold for specific binding (i.e. *k_d_* < 200 nM), showing that for conserved AW-box sites the WRI1 binding affinity tends to be conserved as well (Fig. 4E).

**Figure 4.**
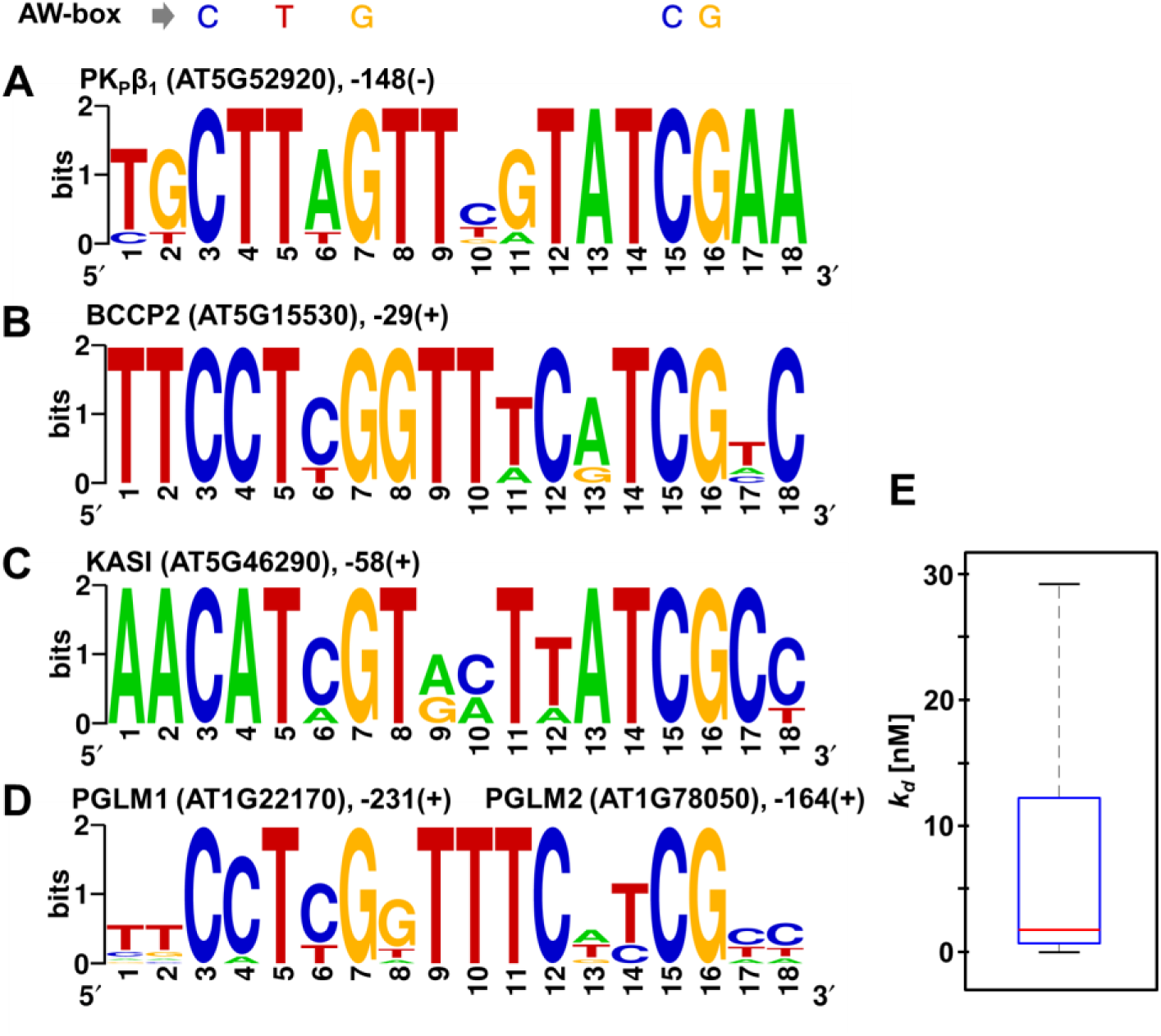
Phylogenetic and functional conservation of AW-box sites found in *A. thaliana* upstream regions. Sequence logos are shown for 5 *A. thaliana* sites that are conserved (upstream positions and directionalities in *A. thaliana* are indicated). A to C, AW-sites conserved within the 12 *Brassicaceae* species of this study. Analyzed by MST were 7 sequences for plastidic pyruvate kinase β_1_-subunit (A), 5 sequences for Biotin Carboxyl Carrier Protein 2 (B), 5 sequences for Ketoacyl-ACP Synthase 1 (C). D, 12 sequences for AW-sites in 2,3-bisphosphoglycerate-dependent phosphoglycerate mutase (PGLM1, PGLM2) conserved within *Brassicaceae* species and outside were analyzed (see also Supplemental Table S7, Supplemental Fig. S6, 7). E, Box-plot showing the total range (black), interquartile range (blue) and median (red) of all 29 *k_d_* values obtained for analysis of DNA fragments in A to D. Aligned sequences and *k_d_* values can be found in Supplemental Table S5. Sequence logos were generated with WebLogo (Crooks et al., 2004).

While exploration of conservation of AW-box sites in this study is focused on the phylogenetic range of *Brassicaceae*, the AW-box can also be shown to be conserved within the flowering plants, as demonstrated for plastidic phosphoglycerate mutase in Fig. 4D. The *A. thaliana* genome contains two chloroplast isoforms for 2,3-bisphosphoglycerate-dependent phosphoglycerate mutase (PGLM1, PGLM2 (Andriotis et al., 2010a). Based to the GenomicusPlants web resource (Louis et al., 2015), the two *A. thaliana* genes result from whole genome duplication events during *Brassicaceae* evolution. Accordingly, Supplemental Table S7 documents collinear/syntenic gene arrangement for the two syntelog *At*PGLM genes. The alignment is extended by genomic regions from 2 Brassicaceae species, 7 dicot species outside the *Brassicaceae* and one monocot species (Supplemental Table S7). Supplemental Fig. S6 documents the high similarity in protein sequences among the PGLM orthologs described in Supplemental Table S7. Close inspection of the genomic regions upstream ATG for the 12 identified orthologous PGLM genes revealed a conserved AW-box motif (Fig. 4D) and all of the aligned sequences showed specific binding activity. A more extensive sequence alignment of PGLM upstream AW-box regions from 19 species is shown in Supplemental Fig. S7. Similar cases of deeply conserved AW-sites for other genes are shown in Supplemental Fig. S8 and S9.

### AW-boxes that bind WRI1 locate close to the transcriptional start site and associated genes tend to be co-expressed with WRI1

It has been shown for several organisms, including *A. thaliana*, that authentic cis-regulatory elements tend to be localized in proximity of the TSS (Yu et al., 2016). Accordingly, Fig. 5A shows that the 62 AW-sites that were identified to specifically bind WRI1 *in-vitro* (*K_d_* < 200 nM, Fig. 2) are positioned close to the TSS. In addition to positional preference, it was assessed whether the genes for which *in-vitro* binding AW-sites were identified are co-expressed with WRI1. WRI1 is mainly expressed during seed development with a characteristic bell-shaped expression pattern (Ruuska et al., 2002). The expression pattern of targets that follow the expression of WRI1 should therefore be positively correlated. Fig. 5B shows that, during seed development, the expression of the 53 gene targets for which specific *in-vitro* binding of AW-boxes to WRI1 was found (*K_d_* < 200 nM) tends to be positively correlated with the expression of WRI1.

**Figure 5.**
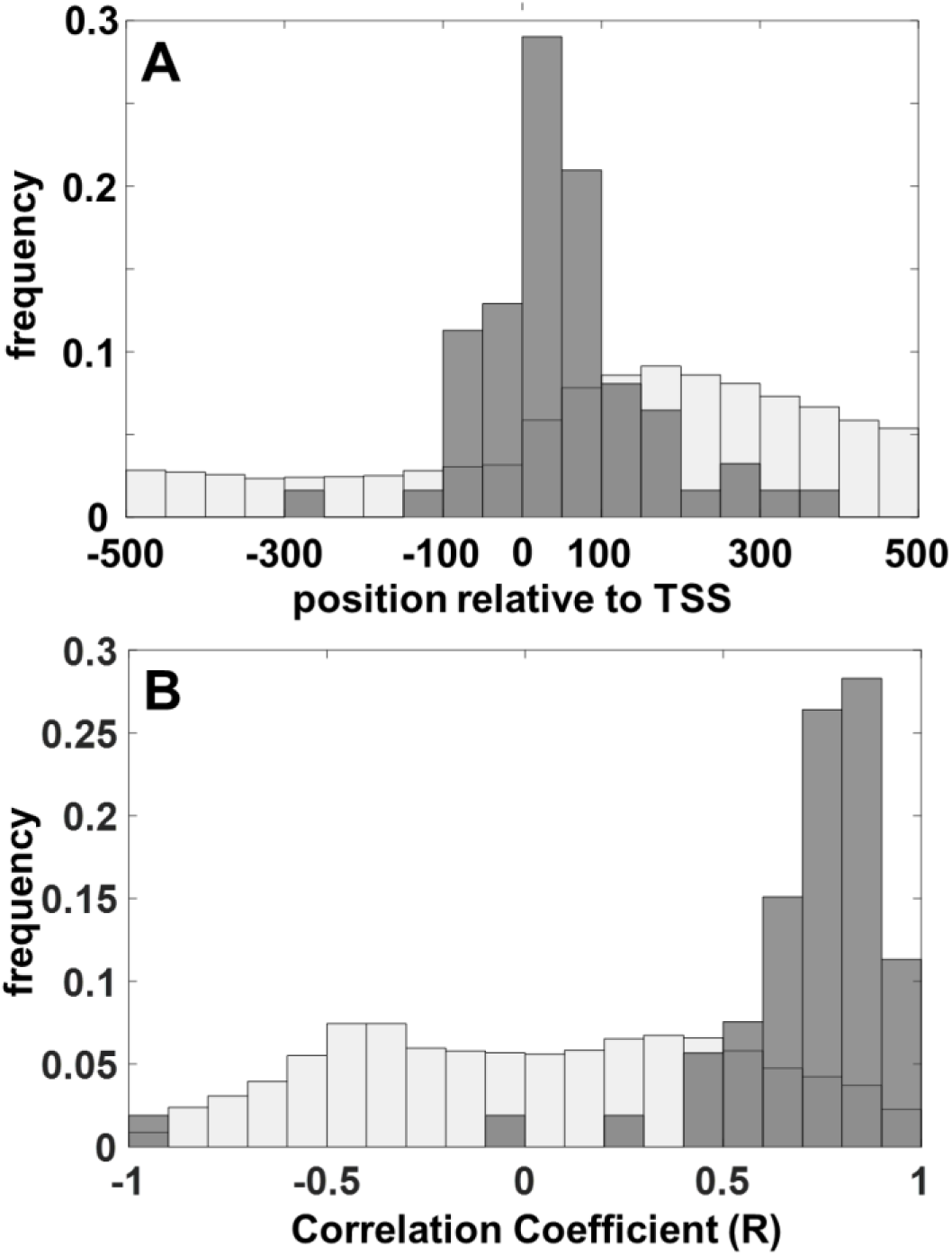
Positional bias of AW-box sites that bind WRI1 *in-vitro* and co-expression analysis of associated genes. A, Positional distribution of WRI1 binding AW-boxes relative to the transcriptional start site (TSS) in *A. thaliana*. Grey bars: 62 AW-box sites identified to be binding WRI1 (*k_d_* < 200 nM, see Fig. 2). White bars: genomic background. B, Co-expression of *A. thaliana* genes with WRI1 during seed development. Pearson Correlation coefficients are based on gene expression data for 8 seed developmental stages taken from the AtGenExpress dataset (Schmid et al., 2005). Grey bars: correlation coefficients for 53 genes for which at least one AW-box can bind WRI1 (*k_d_* < 200 nM, Fig. 2). White bars: genomic background.

### Conserved WRI1 gene targets map to central metabolism

Overall, the motif and gene enrichment workflow shown in Fig. 1 identified 53 *A. thaliana* gene loci as putative WRI1 gene targets. For each gene at least one AW-box site has been characterized as specific *in-vitro* binding to WRI1 and to be conserved (Supplemental Table S5). Examples of this set of genes are highlighted in Table 4 (Sequence logos in Supplemental Fig. S4). In addition, we examined other genes with relevance to central metabolism, FAS and TAG biosynthesis that might have been missed by the enrichment workflow (Supplemental Table S6). For example, enoyl-CoA reductase (ENR) and Ketoacyl-ACP Synthase II (KASII) both are core components of FAS and single copy genes in *A. thaliana*, but no AW-sites were found by the workflow. Manual inspection of genomic sequences revealed AW-box sites positioned further upstream the ATG start than the genome wide applied search window of 1 to 500 bp upstream the ATG start. These sites were shown to specifically bind WRI1 in the *in-vitro* assay and to be well conserved (Supplemental Table S6). With this manual curation of two genes, conserved *in-vitro* binding AW-box sites are found for all steps in FAS between pyruvate kinase and Acyl-ACP thioesterase, the step that releases free fatty acids (Supplemental Table S10). Also, gene targets of the OPPP were inspected as this pathway is likely to contribute reducing equivalents to FAS. All manually curated cases are listed in Supplemental Table S6 and some are highlighted in Table 5 (Sequence logos in Supplemental Fig. S5). Altogether, 68 genes from Supplemental Table S5 and 16 from Supplemental Table S6 were mapped onto central metabolism (Fig. 6). All reactions and genes shown in the figure are identified in Supplementary Table S9. For 60 of 284 shown genes the AW-box was found enriched across ortholog upstream regions and *in-vitro* binding AW-sites were found (Fig. 6). In 46 of these cases *in-vitro* binding AW-sites are highly conserved, with the species conservation ratio above 0.8.

**Figure 6.**
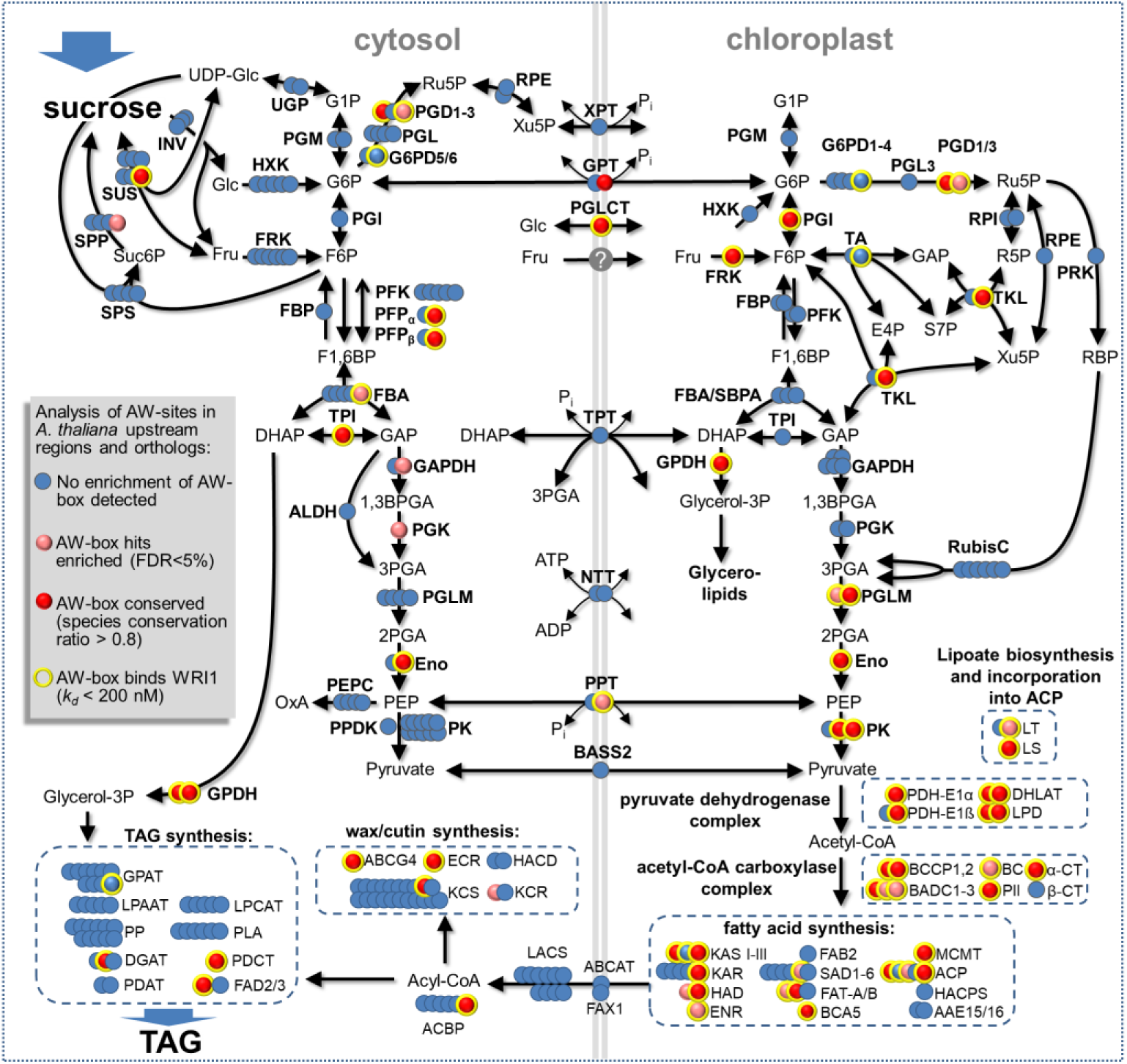
Schematic of carbon allocation in developing seeds of *A. thaliana* with indication of 284 protein encoding genes, highlighting putative WRI1 gene targets with *in-vitro* functional AW-box sites and results on AW-box enrichment and conservation across species. Reactions and genes are identified in Supplemental Table S8. Reaction abbreviations: AAE15/16, Acyl:acyl carrier protein synthetase; ABCAT, ABC Acyl Transporter; ABCG4, ABC Transporter (cutin, wax); ACBP, acyl-CoA-binding protein; ACP, Acyl Carrier Protein; α/β-CT, acetyl-CoA carboxylase carboxyltransferase alpha/beta subunit; ALDH, non-phosphorylating Glyceraldehyde 3-phosphate dehydrogenase; BADC, biotin/lipoyl attachment domain containing; BASS2, Sodium Bile acid symporter family protein; BC, Biotin Carboxylase of Heteromeric ACCase; BCA5, beta carbonic anhydrase 5; BCCP, Biotin Carboxyl Carrier Protein; DGAT, Acyl-CoA : Diacylglycerol Acyltransferase; DHLAT, Dihydrolipoamide Acetyltransferase; ECR, Enoyl-CoA Reductase; Eno, Enolase; ENR, Enoyl-ACP Reductase ; FAB2, Stearoyl-ACP Desaturase; FAD2/3, oleate/linoleate desaturase; FAT, Acyl-ACP Thioesterase; FAX1, Fatty Acid Export 1; FBA, fructose bisphosphate aldolase; FRK, fructokinase; G6PD, glucose 6-phosphate dehydrogenase; GAPDH, Glyceraldehyde 3-phosphate dehydrogenase; GPAT, Glycerol-3-Phosphate Acyltransferase; GPDH, NAD-glycerol-3-phosphate dehydrogenase; HACD, Very-long-chain (3R)-3-hydroxyacyl-CoA dehydratase; HACPS, Holo-ACP Synthase; HAD, Hydroxyacyl-ACP Dehydratase; HXK, hexokinase; INV, invertase; KAR, Ketoacyl-ACP Reductase ; KAS, Ketoacyl-ACP Synthase; KCR, beta-ketoacyl reductase; KCS, 3-ketoacyl-CoA synthase; LACS, Long-Chain Acyl-CoA Synthetase; LPAAT, 1-acylglycerol-3-phosphate acyltransferase; LPCAT, 1-acylglycerol-3-phosphocholine Acyltransferase; LPD, Dihydrolipoamide Dehydrogenase; LS, Lipoate synthase; LT, Lipoyltransferase; MCMT, Malonyl-CoA : ACP Malonyltransferase; NTT, nucleoside triphosphate transporter; PDAT, Phospholipid : Diacylglycerol Acyltransferase; PDCT, Phosphatidylcholine:diacylglycerol cholinephosphotransferase; PDH-E1α, E1-α component of Pyruvate Dehydrogenase Complex; PDH-E1ß, E1-ß component of Pyruvate Dehydrogenase Complex; PEPC, phosphoenolpyruvate carboxylase; PFK, phosphofructokinase; PFP, pyrophosphate dependent phosphofructokinase; PGD, 6-phosphogluconate dehydrogenase; PGI, phosphoglucose isomerase; PGK, Phosphoglycerokinase; PGL, 6-phosphogluconolactonase; PGLCT, plastidic glucose translocator; PGM, phosphoglucomutase; PGL, 6-phosphogluconolactonase; PGLM, phosphoglyceromutase; PII, regulatory subunit of acetyl-CoA carboxylase; PK, pyruvate kinase; PLA, Phospholipase A2; PP, Diacylglycerol-Pyrophosphate Phosphatase; PPDK, pyruvate orthophosphate dikinase; PPT, phosphoenolpyruvate/phosphate antiport; PRK, phosphoribulokinase ; RPE, ribulose-5-phosphate-3-epimerase; RPI, ribose-5-phosphate isomerase; RubisC, ribulose bisphosphate carboxylase; SAD, Stearoyl-ACP desaturase; ß-CT, Carboxyltransferase (Subunit of Heteromeric ACCase); SPP, sucrose-6-phosphate phosphohydrolase; SPS, sucrose phosphate synthase; SUS, sucrose synthase; TKL, transketolase; TPI, Triose phosphate isomerase; TPT, triosephosphate/phosphate antiport; UGP, UDP-glucose pyrophosphorylase; XPT, cylulose 5-phosphate / phosphate translocator. Metabolites abbreviations: 1,3BPGA, ;2PGA, 2-phosphoglyceric acid; 3PGA, 3-phosphoglyceric acid; DHAP, dihydroxyacetone phosphate; E4P, erythrose 4-phosphate; F1,6BP, fructose 1,6 bis-phosphate; F6P, fructose 6-phosphate; Fru, fructose ; G1P, glucose 1-phosphate; G6P, glucose 6-phosphate; GAP, glyceraldehyde 3-phosphate; Glc, glucose; PEP, phosphoenol pyruvate; R5P, ribulose 5-phosphate; RBP, ribulose bisphosphate; Ru5P, ribulose 5-phosphate; Suc6P, sucrose 6-phosphate.

**Table 4.** Examples of putative direct *A. thaliana* WRI1 regulatory gene targets identified by the workflow shown in Fig. 1. Sequence logos are shown in Supplemental Fig. S4. All discovered gene targets, AW-box sequences and further details shown in Supplemental Table S5.

| Gene abbreviation (description, gene ID) | *p*-value for enrichment of AW-box across orthologs | Number of *in-vitro* binding AW-box sites (*A. thaliana*) | WRI1 dissociation constant<sup>1</sup> [nM] | Gene expression correlation with WRI1<sup>2</sup> | species conservation ratio<sup>3</sup> |
| --- | --- | --- | --- | --- | --- |
| ACBP6 (ACYL-COA-BINDING PROTEIN 6, AT1G31812) | 2 x 10<sup>-7</sup> | 1 | 7.9 ± 1.0 | 0.64* | 0.92 |
| BADC1 (Biotin/lipoyl Attachment Domain Containing, AT3G56130) | 2 x 10<sup>-9</sup> | 1 | 0.6 ± 0.2 | 0.69* | 0.83 |
| BADC2 (Biotin/lipoyl Attachment Domain Containing, AT1G52670) | 3 x 10<sup>-6</sup> | 1 | 5.7 ± 2.6 | 0.83** | 0.67 |
| BADC3 (Biotin/lipoyl Attachment Domain Containing, AT3G15690) | 2 x 10<sup>-5</sup> | 1 | 9.5 ± 3.1 | 0.80** | 0.67 |
| BC (Biotin Carboxylase, AT5G35360) | 1 x 10<sup>-5</sup> | 1 | 0.2 ± 0.2 | 0.79** | 0.42 |
| BCA5 (beta carbonic anhydrase 5, AT4G33580) | 5 x 10<sup>-11</sup> | 1 | 0.6 ± 0.5 | 0.72* | 1.00 |
| BCCP1 (Biotin Carboxyl Carrier Protein 1, AT5G16390) | 5 x 10<sup>-9</sup> | 1 | 62.2 ± 19.5 | 0.61 | 0.92 |
| BCCP2 (Biotin Carboxyl Carrier Protein 2, AT5G15530) | 2 x 10<sup>-8</sup> | 2 | 0.7 ± 0.3 | 0.86** | 1.00 |
| bZIP67 (Basic-leucine zipper transcription factor family protein, AT3G44460) | 4 x 10<sup>-9</sup> | 1 | 0.03 ± 0.04 | 0.57 | 1.00 |
| CT-α (Carboxyltransferase α-subunit, AT2G38040) | 1 x 10<sup>-11</sup> | 1 | 70.2 ± 9.4 | 0.86** | 1.00 |
| DGAT2 (Diacylglycerol acyltransferase 2, AT3G51520) | 5 x 10<sup>-9</sup> | 1 | 39.0 ± 9.1 | 0.80** | 0.92 |
| ENO1 (Enolase, AT1G74030)<sup>(p)</sup> | 5 x 10<sup>-10</sup> | 1 | 10.3 ± 6.7 | 0.84** | 0.92 |
| ENO2 (Enolase, AT2G36530)<sup>(c)</sup> | 5 x 10<sup>-6</sup> | 1 | 13.4 ± 3.7 | 0.57 | 0.92 |
| FRK3 (Fructokinase, AT1G66430)<sup>(p)</sup> | 1 x 10<sup>-9</sup> | 1 | 85.2 ± 32.8 | 0.64* | 0.92 |
| GPdHC1 (glycerol-3-phosphate dehydrogenase, AT2G41540)<sup>(c)</sup> | 5 x 10<sup>-11</sup> | 1 | 81.7 ± 15.5 | 0.23 | 0.92 |
| GPdHC1 (glycerol-3-phosphate dehydrogenase, AT3G07690)<sup>(c)</sup> | 7 x 10<sup>-8</sup> | 1 | 105.2 ± 30.8 | 0.79** | 0.83 |
| KASI (Ketoacyl-ACP Synthase I, AT5G46290) | 3 x 10<sup>-11</sup> | 2 | 1.7 ± 1.2 | 0.84** | 1.00 |
| LEC1 (Leafy Cotyledon 1, AT1G21970) | 2 x 10<sup>-7</sup> | 1 | 4.3 ± 1.4 | 0.92** | 0.75 |
| PFP-β (pyrophosphate dependent phospho-fructokinase<sup>(c)</sup> β-subunit, AT1G12000) | 4 x 10<sup>-10</sup> | 1 | 53.9 ± 12.0 | 0.67* | 0.92 |
| PGI1 (phosphoglucose isomerase, AT4G24620)<sup>(p)</sup> | 4 x 10<sup>-5</sup> | 1 | 2.9 ± 0.6 | 0.63* | 0.92 |
| PGLM1 (phosphoglyceromutase, AT1G22170)<sup>(p)</sup> | 7 x 10<sup>-6</sup> | 1 | 1.9 ± 1.3 | 0.91** | 0.75 |
| PGLM2 (phosphoglyceromutase, AT1G78050)<sup>(p)</sup> | 1 x 10<sup>-9</sup> | 2 | 6.7 ± 4.1 | 0.87** | 1.00 |
| PII (PII/GLNB1 homolog, AT4G01900)<sup>(p)</sup> | 2 x 10<sup>-10</sup> | 2 | 9.6 ± 4.1 | 0.83** | 1.00 |
| PK<sub>p</sub>-β<sub>1</sub> (Plastidic pyruvate kinase β<sub>1</sub> subunit, AT5G52920) | 1 x 10<sup>-11</sup> | 2 | 12.2 ± 6.9 | 0.88** | 1.00 |
| PK<sub>p</sub>α (Pyruvate kinase α-subunit, AT3G22960)<sup>(p)</sup> | 1 x 10<sup>-14</sup> | 1 | 0.6 ± 0.3 | 0.89** | 1.00 |
| ROD1 (REDUCED OLEATE DESATURATION 1, AT3G15820) | 3 x 10<sup>-5</sup> | 1 | 2.4 ± 0.3 | 0.70* | 0.92 |
| SUS2 (Sucrose synthase 2, AT5G49190) | 3 x 10<sup>-5</sup> | 1 | 12.6 ± 11.5 | 0.74* | 0.92 |
| TPI (Triose phosphate isomerase, AT3G55440)<sup>(c)</sup> | 2 x 10<sup>-9</sup> | 1 | 76.2 ± 22.2<sup>4</sup> | 0.59 | 0.92 |

<sup>1</sup> For multiple AW-sites per *A. thaliana* gene locus, the lowest measured *k<sub>d</sub>* value is reported here (mean ± SD, *n* = 3). <sup>2</sup>Pearson correlation coefficients for co-expression with WRI1 across 8 seed developmental stages in *A. thaliana* (Schmid et al., 2005). <sup>3</sup> number of species for which an *in-vitro* binding AW-site is conserved / total number of species. <sup>4</sup>consensus “(T/C)CTCGTGATC(AG)TCG” is conserved across 19 sequences (some non-canonical AW-sites). <sup>(c)</sup>Cytosolic compartment isoform; <sup>(p)</sup>plastidic compartment isoform. \*, \*\*Positive correlation (\**p* < 0.05, \*\**p* < 0.01; *n*=8, one-sided test).

**Table 5.** Examples of additional putative direct gene targets of WRI1, examined since being relevant for central carbon metabolism or seed development. Sequence logos are shown in Supplemental Fig. S5. All discovered gene targets, AW-box sequences and further details shown in Supplemental Table S6.

| Gene abbreviation (description, gene ID) | *p*-value for enrichment of AW-box across orthologs | Number of *in-vitro* binding AW-box sites (*A. thaliana*) | WRI1 dissociation constant<sup>1</sup> [nM] | Gene expression correlation with WRI1<sup>2</sup> | species conservation ratio<sup>3</sup> |
| --- | --- | --- | --- | --- | --- |
| AIL6 (AINTÉGUMENTA-like 6, AT5G10510) | $8 \times 10^{-11}$ | 1 | $1.4 \pm 0.8$ | 0.35 | 1.00 |
| CRU3 (cruciferin 3, AT4G28520) | $3 \times 10^{-7}$ | 1 | $1.6 \pm 0.8$ | 0.57 | 0.92 |
| GPD4 (glucose 6-phosphate dehydrogenase, AT1G09420)<sup>(p)</sup> | $2 \times 10^{-3}$ | 1 | $8.2 \pm 4.5$ | 0.87** | 0.50 |
| GPT2 (glucose 6-phosphate/phosphate translocator, AT1G61800) | $5 \times 10^{-5}$ | 1 | $1.6 \pm 0.82^4$ | 0.92** | 0.83 |
| L1L (Leafy Cotyledon1-Like, AT5G47670) | $2 \times 10^{-6}$ | 1 | $10.5 \pm 1.0$ | 0.97** | 0.92 |
| PFP-α (pyrophosphate dependent phosphofructokinase α-subunit, AT1G76550)<sup>(c)</sup> | $3 \times 10^{-1}$ | 1 | $12.3 \pm 4.3^5$ | 0.74* | 0.92 |
| PGD1 (6-phosphogluconate dehydrogenase, AT1G64190)<sup>(c)(p)</sup> | $2 \times 10^{-10}$ | 2 | $9.8 \pm 1.2$ | 0.65* | 0.83 |
| PGD3 (6-phosphogluconate dehydrogenase, AT5G41670)<sup>(c)(p)</sup> | $5 \times 10^{-10}$ | 1 | $9.8 \pm 1.2$ | 0.72* | 0.67 |
| PGLCT (plastidic glucose translocator, AT5G16150) | $2 \times 10^{-10}$ | 1 | $23.9 \pm 4.7$ | 0.75* | 1.00 |
| PPT1 (phosphoenolpyruvate/phosphate translocator, AT5G33320) | $3 \times 10^{-6}$ | 2 | $4.4 \pm 6.6$ | 0.75* | 0.50 |
| TA2 (transaldolase, AT5G13420)<sup>(p)</sup> | $2 \times 10^{-1}$ | 1 | $104 \pm 5.3^6$ | 0.96** | 0.92 |
| TKL1 (transketolase 1, AT3G60750)<sup>(p)</sup> | $3 \times 10^{-11}$ | 1 | $1.0 \pm 0.6$ | 0.53 | 1.00 |

<sup>1</sup> For multiple AW-sites per *A. thaliana* gene locus, the lowest measured $k_d$ value is reported here unless indicated otherwise (mean ± SD, $n = 3$ ). <sup>2</sup> Pearson correlation coefficients for co-expression with WRI1 across 8 seed developmental stages in *A. thaliana* (Schmid et al., 2005). <sup>3</sup> number of species for which an *in-vitro* binding AW-site is conserved / total number of species. <sup>4</sup> Binding affinity given for AW-box canonical ortholog binding site in *Thellungiella halophila*. 9 of 14 other homolog AW-sites bind while the non-canonical homolog site in *A. thaliana* did not bind. <sup>5</sup> Consensus GGTGATCGTATCG conserved across 9 species (non-canonical AW-sites GNTNG(N)<sub>7</sub>CG). <sup>6</sup> consensus "(T/C)(T/C)TTGGTTTG(A/G)TCG" is conserved across 21 sequences (some non-canonical AW-sites). <sup>(c)</sup> Cytosolic compartment isoform; <sup>(p)</sup> plastidic compartment isoform. \*, \*\* Positive correlation (\* $p < 0.05$ , \*\* $p < 0.01$ ; $n=8$ , one-sided test).

This study was based on a highly degenerate binding pattern. The quantitative determinations of WRI1 DNA binding affinity in this study allow for defining WRI1 DNA binding specificity, useful for further computational motif searches. Of the 204 *k_d_* values determined by MST (Supplemental Table S4), 97 are less than 25 nM. Supplemental Table S9 lists position specific base frequencies for the resulting consensus and a Sequence logo emerging from these sequences is shown in Supplemental Figure S11.

## DISCUSSION

The transcription factor WRI1 is as a master regulator of lipid accumulation in developing seeds of *A. thaliana* and other plants (Cernac and Benning, 2004; Kong and Ma, 2018). While WRI1 is known to directly transcriptionally activate genes in FAS, we hypothesized that additional enzyme steps for sucrose catabolism between sucrose synthase and pyruvate kinase might also be direct targets. To increase confidence for identification of cis regulatory elements, a comparative genomics approach might be useful. Former studies have indicated that intragenomic comparison between *A. thaliana* and close relatives has the potential to discover a conserved *cis* regulome. For example, by comparative analysis of 9 *Brassicaceae* genomes, Haudry *et al*. (2013) mapped conserved noncoding sequences for different genomic regions and found them to cover close to 10 % of the 5’UTR regions and the regions from 1 to 200 bp upstream the TSS in the *A. thaliana* genome. Such conserved regions tend to be enriched in known regulatory motifs and to be under selective pressure (Haudry et al., 2013). Here we developed a genome-wide analysis workflow to identify *A. thaliana* AW-box sites that are phylogenetically preserved across *Brassicaceae* species, followed by pathway analysis and quantification of WRI1 binding affinity by an *in-vitro* DNA binding assay (Fig. 1). While this process does not provide direct evidence of *in-vivo* functionality, multiple lines of evidence support the utility of our approach: 1) Our workflow identified a number of well-characterized AW-box sites for BCCP2, KASI, PK_p_-ß_1_ and SUS2 (Maeo et al., 2009) (Table 4) along with genes controlling all steps of FAS (Supplemental Table S10). 2) Conservation of the AW-box sequences indicates that they are under purifying selection. For 5 select cases we demonstrated that WRI1 binding activity that was measured for *A. thaliana* AW-sites is retained in phylogenetically conserved orthologous AW-box sites in the other Brassicaceae species investigated (Fig. 4). 3) For 53 *A. thaliana* gene loci identified herein, 62 high-affinity AW-binding sites were located close to the TSS (Fig. 5A), consistent with findings by Fukuda *et al*. (2013), suggesting that the *in-vivo* functionality of AW-box sites strongly depends on their proximity to the TSS. 4) Most of the 53 genes associated with the *in-vitro* AW-sites binding are co-expressed with WRI1 during seed development (Fig. 5B).

Fig. 6 graphically illustrates the information regarding AW-box enrichment across orthologous upstream regions in addition to WRI1 binding affinity onto a metabolic pathway scheme related to seed oil synthesis. For the conversion of pyruvate into fatty acids, evidence for AW-box site conservation and tight WRI1 binding affinity is found for 27 genes in three major protein complexes (Fig. 6). These include the chloroplast pyruvate dehydrogenase complex, acetyl-CoA carboxylase and fatty acid synthesis. Besides the canonical components of FAS, Beta Carbonic Anhydrase 5 is a likely WRI1 target (BCA5, Table 4) and might therefore be of general relevance for FAS. *At*BCA5 has been shown to be targeted to the chloroplast (Fabre et al., 2007). We propose that the enzyme might be required for conversion of CO_2_ to bicarbonate (HCO_3_ ^-^) when FAS operates at high rates. This is because within FAS, acetyl-CoA carboxylase (ACC) and ketoacyl-ACP synthase (KAS) create a cycle of carboxylation and decarboxylation where ACC requires bicarbonate (HCO_3_ ^-^) as a substrate (Li-Beisson et al., 2013) while KAS releases CO_2_. In support of a requirement for carbonic anhydrase for high FAS rates, specific BCA inhibitors have been shown to inhibit FAS in developing embryos of cotton (*Gossypium hirsutum*) (Hoang and Chapman, 2002).

Similar to the concerted control of most genes encoding FAS enzymes by WRI1, a contiguous lower section of glycolysis is recognized as being controlled by WRI1 (Fig. 6; Table 4, 5), including the plastidic phosphoglycerate mutase (PGLM1, PGLM2) and two subunits of plastidic pyruvate kinase (PK), plastidic and cytosolic enolase (Eno) as well as the phosphoenolpyruvate/phosphate translocator (PPT1). A similar coherent pathway section can be recognized in pyrophosphate dependent phosphofructokinase (PFP), the cytosolic isoforms of fructose bisphosphate aldolase (FBA), triose phosphate isomerase (TPI) and NAD-glycerol-3-phosphate dehydrogenase (GPDH) (Fig. 6). In other pathway sections seen in the upper half of glycolysis and the PPP evidence for WRI1 control is more sparsely distributed. However, some new gene targets in the upper section of the pathway yield some insights on possible modes of functioning of this complex and redundant network during seed oil deposition. For example, sucrose is cleaved by invertase or sucrose synthase. In either case fructose is obtained, which in turn can be transformed by hexokinase (HXK) or fructokinase (FRK) into fructose 6-phosphate (F6P) (Fig. 6). Among 5 HXK and 7 FRK genes identified by our motif and gene enrichment workflow, only FRK3 was identified as a likely WRI1 target (Table 4, chloroplast localized FRK in Fig. 6). This finding can be rationalized with respect to a recent complete biochemical and genetic characterization of the FRK gene family in *A. thaliana* (Stein et al., 2016; Riggs et al., 2017). While no severe seed phenotype was found for the single-KO mutation of FRK3, a FRK1-FRK3 double-KO mutation resulted in a severe wrinkled seed phenotype with strong reduction in seed oil content (Stein et al., 2016). In conclusion, while our approach singled out FRK isoform 3 as a new WRI1 target, the genetic evidence shows that seed oil synthesis depends predominantly, although not exclusively, on contributions of this specific isoform. In addition to FRK3 we identified plastidic phosphoglucose isomerase (PGI1) as a putative WRI1 target (Table 4, Fig. 6). As in the case of FRK3, there is recent genetic evidence in support of PGI1 being important for TAG synthesis. A PGI1 mutant (*pgi1-2*) was reported to have reduced seed yield per plant, seed size and seed oil content (Bahaji et al., 2018). Reciprocal crosses of *pgi1-2* with wild type showed that the low oil, wrinkled seed phenotype is independent of maternal influences (Bahaji et al., 2018). This strongly corroborates our finding of PGI1 being a WRI1 target and thus important for oil synthesis.

Additional putative WRI targets relate to the OPPP, which is considered a major source of reductant in heterotrophic plant tissues, delivering NADPH for various biosynthetic processes (Neuhaus and Emes, 2000; Kruger and von Schaewen, 2003), including FAS (Neuhaus and Emes, 2000; Rawsthorne, 2002). We identified several OPPP-associated genes as likely WRI1 targets (see Table 5, Fig. 6): Transketolase (TKL1), transaldolase (TA2), and two isoforms of 6-phosphogluconate dehydrogenase PGD3 and PGD1, the two of which accumulate both in the cytosol and the chloroplast (Holscher et al., 2016). Based to the GenomicusPlants web resource (Louis et al., 2015), PGD1 and PGD3 result from a whole genome duplication event at the basis of the *Brassicaceae*. One AW-box site appears to be conserved for both genes (Supplemental Table S6) and is also found more widely conserved across dicot species (Supplemental Fig. S9). WRI1 binding assays also implicate functional AW-sites for isoforms of glucose 6-phosphate dehydrogenase, but less well conserved (G6PD4, Table 5).

During seed storage synthesis, chloroplast biosynthetic activities might depend on the movement of hexose carbon between cytosol and chloroplast. Our analysis implicates related transport functions to be controlled by WRI1 (Fig. 6). Previous literature has emphasized the role of the glucose 6-phosphate/phosphate translocator (GPT) for entry of glucose 6-phosphate into the chloroplast during early phases of seed development in *A. thaliana* and *B. napus* (Eastmond and Rawsthorne, 2000; Ruuska et al., 2002; Kubis et al., 2004). *A. thaliana* contains two GPT genes. AtGPT1 (AT5G54800) is reported to be essential in pollen and early embryo development (Niewiadomski et al., 2005; Andriotis et al., 2010b), while loss of AtGPT2 (AT1G61800) seems not to have a severe phenotype (Niewiadomski et al., 2005). AtGPT1 has been ascribed a role in carbon import during early and later stages of seed development when it may be relevant to feed into starch synthesis and FAS (Ruuska et al., 2002; Niewiadomski et al., 2005; Andriotis et al., 2010c). Here we found a conserved binding site in the promoter regions of GPT2 orthologs (Table 5) and some of the orthologous sites were found to bind WRI1 *in-vivo*, hinting at functional relevance of GPT2 in oil synthesis. However, the sites in GPT2 of *A. thaliana* and three other species are divergent from the AW-box consensus and did not bind in the *in-vitro* WRI1 binding assay (Supplemental Table S6), making it more ambiguous as to whether GPT2 is controlled by WRI1. In addition to GPT2, we found a conserved AW-box in the promoter of the plastidic glucose translocator (PGLCT) (Weber et al., 2000) (Table 5, Fig. 6). The PGLCT has been implicated to be involved in photo-assimilate export from leaf chloroplasts when starch mobilization takes place in the dark (Cho et al., 2011). If this transporter has a role in a heterotrophic context in seed development during oil accumulation, it likely facilitates entry of glucose into the chloroplast rather than export. In support of this idea, isolated chloroplasts of *Brassica napus* developing embryos have been shown to have the ability to take up glucose with a saturation kinetics consistent with transporter facilitated uptake (Eastmond and Rawsthorne, 1998). Moreover, the uptake capacity of embryo chloroplasts was substantially higher than that of isolated leaf chloroplasts (Eastmond and Rawsthorne, 1998).

In addition to the importance of transport of hexose phosphates or hexose across the chloroplast envelope for oil synthesis, the cytosolic glycolysis pathway has been deemed to be important for seed filling as well (White et al., 2000; Ruuska et al., 2002). More recent metabolic flux studies in *B. napus* developing embryos give further support to substantial carbon flux through cytosolic glycolysis with particular relevance of the phosphofructokinase reaction in flux control (Schwender et al., 2015). In the cytosol, pyrophosphate-fructose-6-phosphate-phosphotransferase (PFP, EC 2.7.1.90) is an allosterically controlled heteromeric complex of α-and β-subunits (Mustroph et al., 2013). During seed development in different oilseed species, PFP subunits seem to be expressed at higher levels than the cytosolic ATP-dependent PFK (Troncoso-Ponce et al., 2011). Consistent with the suspected relevance of cytosolic PFP for seed oil synthesis we found conserved *in-vitro* functional AW boxes in the upstream regions of both α- and -β subunits of PFP (Table 4, 5) (Fig. 6).

Although TAG biosynthesis genes are not generally viewed as direct targets of WRI1, our data identify several candidates for direct WRI1 targeting associated to the TAG biosynthetic sub-network (Fig. 6): These include two isoforms of cytosolic glycerol 3-phosphate dehydrogenase (GPDH), Acyl-CoA binding protein 6 (ACBP6), Diacylglycerol acyltransferase 2 (DGAT2) and REDUCED OLEATE DESATURATION1 (ROD1) (Table 4, Fig. 6). Of the three different and functionally non-redundant types of DGAT found in plants, DGAT1 has been suggested to carry most of the TAG synthesis during seed development (Li-Beisson et al., 2013). However, our results suggest that DGAT2 is under direct control of WRI1 (Table 4). In contrast to DGAT1, *At*DGAT2 has been reported to have preference for 18:3-CoA relative to other fatty acid CoA esters (Zhou et al., 2013), which could point to a contribution of DGAT2 to channeling polyunsaturated fatty acids into TAG. One of the other genes putatively under direct control of WRI1, ROD1, encodes for phosphatidylcholine:diacylglycerol cholinephosphotransferase (PDCT) and has also been implicated in regulating the poly-unsaturation state of TAG (Lu et al., 2009). Consistent with WRI1 having control over ROD1 expression, ROD1 expression is significantly increased when WRI1 is overexpressed in *A. thaliana* developing seeds (Adhikari et al., 2016) or in *Nicotiana benthamiana* leaves (Grimberg et al., 2015). In addition, ROD1 expression was found to be reduced in the *wri1 wri3 wri4* mutant (To et al., 2012). Another TAG biosynthetic enzyme putatively under direct control of WRI1 and involved in TAG synthesis is Glycerol 3-phosphate dehydrogenase, which provides the glycerol backbone for TAG. In *A. thaliana*, two cytosolic isoforms (AT2G41540, AT3G07690) have been identified (Shen et al., 2006) as well as one plastidic one AT5G40610 (Wei et al., 2001). Remarkably, we have found indications for control by WRI1 for all three genes (Fig. 6).

The transcription factor WRI1 is positioned towards the end of a gene regulatory cascade governing seed development and storage accumulation. WRI1 is likely under direct control of LEAFY COTYLEDON1 (LEC1), LEAFY COTYLEDON2 (LEC2) and FUSCA3 (FUS3) (Fatihi et al., 2016). Data presented herein suggests that WRI1 directly controls its own regulator, LEAFY COTYLEDON1 (Table 4), implying a condition of positive or negative autoregulation. In addition, LEAFY COTYLEDON1-LIKE (L1L) (Table 5) as well as the basic leucine zipper transcription factor 67 (bZIP67) (Table 4) were found as putative WRI1 targets. L1L is known to act in association with bZIP67 in transcriptional activation of several genes related to seed storage accumulation (Yamamoto et al., 2009; Mendes et al., 2013), which includes cruciferin 3 (CRU3) and SUS2 (Yamamoto et al., 2009) for both of which we found conserved high affinity AW-boxes (Table 4, Table 5).

### Conclusions

The motif and gene enrichment workflow applied in this study identified numerous known WRI1 targets related to oil synthesis in addition to a battery of additional genes, including those coding for enzymes for the conversion of sugars to pyruvate and enzymes in the TAG biosynthesis sub-network. While this study was mostly limited to the *Brassicaceae* family, we provided exploratory examples of AW-sites being deeply conserved in other plant families (Supplemental Fig. S7-S9). In this work we demonstrate that a workflow based on the identification of phylogenetically conserved binding sites along with highly quantitative *in-vitro* binding assays can be a powerful approach to identify potential regulatory networks. The analysis presented herein can be generally applied to other TFs and expanded to other plant families.

The GO term and pathway analysis stage of the genome analysis workflow (Fig. 1) narrowed down the initial enriched gene set to 73 putative WRI1 targets that belong to specific pathways in seed oil synthesis. It is likely that there are additional targets of WRI1 that are outside of oil biosynthesis (Kong et al., 2017; Liu et al., 2019), and that it might be important to explore these if WRI1 is to be used for metabolic engineering of oil production (Kong et al., 2019). Such additional targets might have been excluded in the pathway enrichment step of our process. Further refinements of the comparative genomics approach presented herein might be helpful to identify such targets. Another aspect of functional overlap not explored here is cuticular wax biosynthesis. It has been shown that *A. thaliana* WRINKLED4, a close homolog to WRI1, regulates cuticular wax biosynthesis and has many gene targets in common with WRI1 (Park et al., 2016). Since Fig. 6 shows three WRI1 gene targets related to wax/cutin synthesis it would be interesting to use the same approach to evaluate WRINKLED4 gene targets and investigate its regulatory overlap with WRI1.

## MATERIAL AND METHODS

### Data sources and processing

Sequence files corresponding to upstream and downstream sequences of the annotated start codon/end codon of *Arabidopsis thaliana* were downloaded from The Arabidopsis Information Resource (TAIR) (Rhee et al., 2003), version 10 (See details in Supplemental Table S1). For 12 Brassicaceae species used in this study, files for genome sequences, protein sequences and genome annotation (General Feature Format, GFF) were retrieved from sources listed in Supplemental Table S1. In-house generated PERL and PYTHON scripts were used to process and analyze the sequence information. Data processing and analysis was also done using Microsoft Excel (http://www.microsoft.com) and with Matlab (version R2016a, The MathWorks, Inc., Natick, Massachusetts, United States).

Gene annotation and classification information on *Arabidopsis thaliana* lipid metabolism genes in was collected from the ARALIP database (http://aralip.plantbiology.msu.edu/data/aralip_data.xlsx, accessed June 29, 2017) (Li-Beisson et al., 2013; McGlew et al., 2014). The database contains an expert curated list of 822 reactions / proteins associated to lipid metabolism. 775 AGI gene locus identifiers are associated to 24 lipid pathways. In addition, we used a pathway classification as given by *Bna572*, a manual curated large-scale metabolic model for the storage compound accumulation during seed development in *Brassica napus* (Hay et al., 2014). 962 AGI gene locus identifiers are categorized into 77 metabolic pathways. As a more comprehensive source we used MapMan hierarchical classifications for *Arabidopsis* genes (http://www.gabipd.org/database/java-bin/MappingDownloader)(Usadel et al., 2009).

### Synteny analyses for Brassicaceae genomes and other species

Syntenic orthology relations between protein encoding genes of *A. thaliana* and other species were derived by using the SynOrths tool (version 1.0, Cheng et al., 2012). In short, this tool derives pairwise synteny relations based on protein sequence similarity and gene adjacencies on contigs (Cheng et al., 2012). For each comparison, the *A. thaliana* genome was always defined as the target genome. Default parameter settings were applied since they have already been optimized for the closely related Brassicaceae genomes (Cheng et al., 2012). In addition to SynOrths outputs, published synteny information related to *Brassica napus* and *Camelina sativa* was used as additional reference (Chalhoub et al., 2014; Kagale et al., 2014). Within the set of Brassicaceae genomes used here, the genome assembly of *Aethionema arabicum* seemed to be of lesser quality, as judged by the size distribution of contig lengths (Haudry et al., 2013). Therefore, homology relations between the *A. thaliana* genome and *A. arabicum* was derived only based on similarity between sequences of predicted proteins. Protein sequence alignments were established by BLAST (protein sequence similarity) (Altschul et al., 1997) with the predicted protein sequences of *A. arabicum* as query against a database of TAIR10 predicted proteins (representative gene models). BlastP results were filtered with an E-value cut-off of 10^-7^. Using a custom script, alignments spreading over multiple lines (broken alignments) were joined. Top hits were retained as homology relations but rejected if the alignment length was less than 70 % of the length of the query, or if the percentage of identical matches was below 60 %.

Besides the automated analysis, syntenic relationships were tracked manually in a few cases to explore phylogenetic conservation from *A. thaliana* to species outside the Brassicaceae. Genomic sequences of *Citrus sinensis*, *Daucus carota*, *Populus trichocarpa*, *Ricinus communis*, *Sesamum indicum*, *Sorghum bicolor*, *Tarenaya hassleriana* and *Vitis vinifera* were mined using the online resources of the National Center for Biotechnology Information (NCBI) (www.ncbi.nlm.nih.gov) (Coordinators, 2018). Protein searches among the species was done using NCBI BLAST (Johnson et al., 2008). To identify the gene order in chromosomal neighborhoods of genes of interest of the non-Brassicaceae species, genomic information (GFF files) was accessed at NCBI from sources as listed in Supplemental Table S1.

### Mining of genomic DNA sequences

Nucleotide sequences 500 bp upstream of the ATG start codon were extracted for each protein encoding gene locus and for each genome of the 12 Brassicaceae species. For this purpose, GFF files (Supplemental Table S1) were mined for genomic coordinates of start codons. The tool gffread (Trapnell et al., 2010) was then used to extract DNA sequences from -1 to -500 nt upstream the start codon from genomic sequences. In case of *A. thaliana*, sequences were extracted only for one gene model per locus (representative gene models). In particular, for each gene in the sequence file “TAIR10_upstream_500_translation_start_20101028” (Supplemental Table S1) start codon positions were identified from sequence headers and matched to one gene model version in the TAIR10 gff file (genomic feature “protein”). This allowed to define genomic coordinates for regions upstream and downstream the transcriptional start codon and downstream the stop codon for each representative gene model.

### Search of DNA sequences for string pattern matches

Searching DNA sequences for all occurrences of a string pattern was done using an in-house tool. The search tool was tested for accuracy by comparing search outputs with the outputs of another pattern search tool (http://www.bioinformatics.org/sms2/dna_pattern.html)(Stothard, 2000). For each *A. thaliana* gene locus different genomic regions (Upstream and downstream ATG start, intron sequences, downstream stop codon) were searched for occurrence of the AW-box (5’-CNTNG(N)_7_CG-3’, N = A, C, T or G) in both sense and antisense directions. Overlapping motif hits were recognized. For further processing, detected motif matches were recorded along with the gene ID and genomic position of the searched sequence as well as the position and orientation of detected sequences relative to the ATG start.

### Collection of AW-box sites and assessment of sequence conservation

Nucleotide sequences 500 bp upstream of the ATG start codon from 12 Brassicaceae species were searched for the AW-box binding consensus in both strands and matches were recorded with adjacent sequence context as 18 nc sequences (5’-NNCNTNG(N)_7_CGNN-3’). To trace conserved motif instances, AW-box sites found in *A. thaliana* upstream regions were compared to sites found in OURs (pairwise un-gapped alignments). We considered sequence conservation to be given if two motif instances were identical in orientation relatively to the ATG start and if the sequence comparison of the two 18 nc sequences had at least 14 identities (77.8 % identity). This identity threshold was empirically derived by comparison of randomly generated AW-box sites, given a 33.2 % G+C content as found in upstream regions of *A. thaliana* (Supplemental Fig. S3A). The mean expectation of identities between two random sampled AW-box sites was 8.613 and 14 or more identities were obtained for 0.2 % of comparisons, i. e. the by-chance probability to judge two random AW-sites to be conserved is 0.002 (Supplemental Fig. S3A). Conserved sequences from multiple pairwise comparisons between an *A. thaliana* AW-box site and orthologous sites were aggregated into sets of conserved sequences and further into sets of conserved species (Supplemental Fig. S3B). The number of conserved species divided by 12 (total number of assessed species) is the species conservation ratio. If AW-box site binding affinity to WRI1 protein was measured, the species conservation ratio was determined by only tracking conservation of specific binding AW-sites.

### Sequence logos and computation of information content

Sequence logos were created using the WebLogo version 2.8.2 online tool (Crooks et al., 2004)(http://weblogo.berkeley.edu/). The logos created in this study are intended to compare alignments of binding site sequences between each other, not to quantify how degenerate a site is relative to a genomic background. This is important because WebLogo assumes uniform background symbol distribution (all four bases appear with equal background frequency of 0.25), while the genomes studied here have significantly skewed background base distributions. To compare sequence alignments based on numerical values, the information content was calculated as described by Workman *et al*. (2005), based on a uniform background base composition and using the logarithm base 2. For computation of base frequencies when using small datasets, a pseudo-count value of 0.0001 was used.

### Enrichment analysis of presence of AW-box motifs in different Arabidopsis genome features

To model the occurrence of a motif in gene regions of uniform size (e.g. 500 bp upstream the ATG start codon), the hypergeometric distribution was applied. One or more matches of the AW-box motif were counted as a motif hit. If among a total population *N* (total number of searched gene regions) there are *K* motif hits, then the probability to find *m* motif hits in a sub-set of *n* searched gene regions is given by the probability mass function for the hypergeometric distribution:

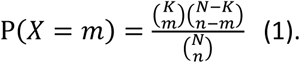

To assess the expected value for *m* AW-motif hits among the *n* = 52 FAS genes, P(*X*=*m*) was computed for *m* ranging within zero and 52 by using the respective Microsoft EXCEL® spreadsheet function (HYPGEOM.DIST). From the results the approximate range of *m* for which 99.9 % of the observations are to be expected (99.9 % confidence interval) was determined symmetrically around the mean expectation value for *m* (*nK*/*N*).

To test for enrichment of the AW-box across sets of OURs, the cumulative hypergeometric probability mass function was applied. Here *N* is the number of protein-encoding genes for which 500 bp upstream regions were searched among all 12 genomes and *K* is the total number of AW-box hits. In case of overrepresentation (*m* ≥ *n*\**K*/*N*), the probability of observing *m* or more AW-box hits within a sample of *n* genes (ortholog group size) is given by:

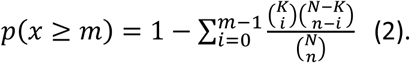

The false positive rate was estimated with an empirical null model: All *p*-value computations were repeated after randomization of the upstream sequences, using the tool “fasta-shuffle-letters” from the MEME suite (Bailey et al., 2009). For a given significance threshold, *t*, the number of significance calls for the randomized sequence data (*p*-value ≤ *t*), divided by the number of significance calls for the unperturbed sequence data was taken to be the False Discovery Rate (FDR).

To test for enrichment of the AW-box in *A. thaliana* intron sequences, the sequence file “TAIR10_intron_20101028” was searched. Since the searched DNA sequences were not of uniform size, the hypergeometrical model was not applied. An estimation for the frequency of motif matches by chance was done based on random shuffling of the intron sequences for the 52 FAS genes.

### Pathway and GO-term enrichment analysis

The test for gene enrichment in pathways was done using equation 2 to assess the probability of observing *m* or more *A. thaliana* genes for which the AW-box is conserved across OURs within a set of *n* genes representing a pathway. In this case *N* is the number of ortholog gene sets and *K* the number of *A. thaliana* genes for which the AW-box was significantly enriched across OURs. Only pathway genes that are part of the background set *N* were counted. Adjustment for multiple hypothesis testing was done by multiplying the *p*-values with the total number of gene sets tested for (Bonferroni correction). Only gene sets with 2 or more members were counted.

Gene Ontology (GO) term enrichment analysis for sets of Arabidopsis genes (TAIR ID) was performed with the functional annotation tool of the online bioinformatics resources given by the Database for Annotation, Visualization and Integrated Discovery (DAVID) (v6.8; https://david.ncifcrf.gov/, Jiao *et al*.,(2012)).

### Evaluation of publicly accessible gene expression data

To analyze gene expression during seed development we re-analyzed microarray gene expression data from Schmidt et al., (2005). In short, *Arabidopsis* developmental series microarray gene expression data with 22,810 signals for 79 developmental stages and different tissues in plant development were analyzed (http://www.ncbi.nlm.nih.gov/bioproject/96937). Expression values for 8 stages of seed development (sample identifiers ATGE_76 to ATGE_84) were averaged across the three replicates. Pairwise Pearson’s correlation coefficients were computed between WRI1 (At3g54320, Affymetrix identifier “251891_at”) and each of the other array signals. To disregard lower intensity expression values, correlation values were only derived for 17058 signals, which is the upper 75^th^ percentile of the sum of expression values across the 8 seed development stages. Significance of Pearson’s correlation coefficients (sample size *n*) was tested for with the Student’s *t*-test for *n*-2 degrees of freedom with critical *t-*values derived from correlation coefficients as described by Olson (1987).

### Microscale thermophoresis

Specific binding of AW sites by AtWRI1 was measured by MST (Seidel et al., 2013). A genetic construct combining the coding region responsible for DNA binding, green fluorescent protein (GFP) and a HIS-tag (*At*Wri_150-240_-GFP-HIS) was expressed in *E. coli* and the protein purified as described before (Liu et al., 2019). Thermophoretic assays were conducted using a Monolith NT.115 apparatus (NanoTemperTechnologies, South San Francisco, CA; nanotempertech.com). Assay conditions were as previously (Liu et al., 2019). dsDNA oligomers were hybridized using a thermal cycler. For determining a dissociation constant (*k_d_*), 16 reactions were prepared: 8 nM of-AtWRI1_58-240_-GFP was incubated with a serial (1:1) dilution of the ligand (dsDNA) from 1.25 µM to 38.81 pM or from 6.25 µM to 190 pM. Samples of approximately 10 μl were loaded into capillaries and inserted into the MST NT.115 instrument loading tray. All thermophoresis experiments were carried out at 25 °C using 40% MST power and 100% or 80% LED power. The data were fitted with the NanoTemper Analysis software v2.2.4. To ensure reproducibility, any series of measurements performed at one day included a reference DNA probe (Ligand 10, Supplementary Table S4). For this ligand the average binding affinity (*K_d_*) out of multiple series of measurements was 7.0 ± 3.5 nM. In each series of measurements, *k_d_* values given by the analysis software were only accepted if the response amplitude was within about ± 20 % of the response amplitude measured for the reference DNA probe (Supplemental Fig. S10). Otherwise, the DNA fragmented was assessed to be “not binding”. For examples of evaluation of analysis MST results see Supplemental Fig. S10. Statistics for *k_d_* values (standard deviation) were derived from three measurements of a DNA ligand obtained from different measurement series.

## Accession Numbers

Sequence data from this article can be found in the Arabidopsis Genome Initiative or GenBank/EMBL databases under the following accession numbers: ACBP6(AT1G31812), AIL6(AT5G10510), BADC1(AT3G56130), BADC2(AT1G52670), BADC3(AT3G15690), BC(AT5G35360), BCA5(AT4G33580), BCCP1(AT5G16390), BCCP2(AT5G15530), bZIP67(AT3G44460), CRU3(AT4G28520), CT-a(AT2G38040), DGAT2(AT3G51520), ENO1(AT1G74030), FRK1(AT5G51830), FRK3(AT1G66430), G6PD4(AT1G09420), GPDHc1(AT2G41540), GPDHc2(AT3G07690), GPDHp(AT5G40610), GPT1(AT5G54800), GPT2(AT1G61800), KASI(AT5G46290), L1L(AT5G47670), LEC1(AT1G21970), LOS2(AT2G36530), PFP-a(AT1G76550), PFP-ß(AT1G12000), PGD1(AT1G64190), PGD3(AT5G41670), PGI1(AT4G24620), PGLCT(AT5G16150), PGLM1(AT1G22170), PGLM2(AT1G78050), PII(AT4G01900), PKp-α(AT3G22960), PKp-ß1(AT5G52920), PPT1(AT5G33320), ROD1(At3g15820), SUS2(AT5G49190), TA2(AT5G13420), TKL1(AT3G60750), WRI1(AT3G54320).

## Supplemental Data

**Supplemental Table S1**: Sources for genomic information.

**Supplemental Table S2:** Enrichment of the AW-box motif in genomic features of genes encoding for fatty acid biosynthesis in *Arabidopsis thaliana*

**Supplemental Table S3**: GO term and pathway enrichment for 915 *A. thaliana* genes for which the AW-box is significantly enriched in upstream regions across orthologs.

**Supplemental Table S4**: Analysis of binding affinity of DNA fragments to AtWRI1 by Microscale Thermophoresis (MST).

**Supplemental Table S5**: Conservation of the AW-box motif in regions upstream ATG for 12 brassicaceae species (73 genes identified by pathway analysis).

**Supplemental Table S6**: Conservation of the AW-box motif in regions upstream ATG for 12 brassicaceae species (21 additional genes).

**Supplemental Table S7**: Syntenic gene analysis for genomic regions close to 2,3-bisphosphoglycerate-dependent phosphoglycerate mutase (PGLM) in 11 species.

**Supplemental Table S8**: Identification and details on *Arabidopsis thaliana* genes shown in Figure 6.

**Supplemental Table S9:** Positional Frequency Matrix (PFM) for 97 AW-box sites with measured *k_d_* values of less than 25 nM (MEME format).

**Supplemental Table S10.** AW-box motif found for genes in the *A. thaliana* genome encoding components related to fatty acid biosynthesis.

**Supplemental Figure S1:** Distribution of ortholog gene set size after synteny analysis.

**Supplemental Figure S2.** Enrichment of the AW-box across orthologous upstream regions of 12 *Brassicaceae* genomes.

**Supplemental Figure S3.** Conservation of AW-box sites between *Arabidopsis thaliana* and other species.

**Supplemental Figure S4:** Conserved AW-box sites for WRI1 gene targets discovered by the motif and gene enrichment workflow.

**Supplemental Figure S5:** Conservation of the AW-box sites for additional WRI1 gene targets.

**Supplemental Figure S6:** Alignment of protein sequences for chloroplast 2,3-bisphosphoglycerate-dependent phosphoglycerate mutase from 11 plant species.

**Supplemental Figure S7.** Deep conserved AW-box site in upstream regions of chloroplast isoforms for 2,3- bisphosphoglycerate-dependent phosphoglycerate mutase (PGLM) of mono- and eudicot plants.

**Supplemental Figure S8.** Deep conserved AW-box site in upstream regions of plastidic pyruvate kinase β_1_-subunit of mono- and eudicot plants.

**Supplemental Figure S9.** Deep conserved AW-box site in upstream regions of 6-phosphogluconate dehydrogenase (PGD) genes of eudicot plants.

**Supplemental Figure S10.** Examples for binding curves of AtWRI1_58-240_-GFP and 28 bp DNA ligands.

**Supplemental Figure S11.** Sequence logo for 97 sequences with *k_d_* value < 25 nM.

## ACKNOWLEDGEMENTS

This work was supported by the U.S. Department of Energy, Office of Science, Office of Basic Energy Sciences under contract numbers DE-SC0012704 (to J.Sc.) and KC0304000 (to J.Sh.) - specifically through the Physical Biosciences program of the Chemical Sciences, Geosciences and Biosciences Division.

